# The genetic architecture of quantitative variation in the self-incompatibility response within *Phlox drummondii* (Polemoniaceae)

**DOI:** 10.1101/2025.05.27.656016

**Authors:** Grace A. Burgin, Federico Roda, Matthew Farnitano, Charles Hale, Antonio Serrato-Capuchina, Robin Hopkins

**Affiliations:** Department of Organismic and Evolutionary Biology, The Arnold Arboretum, Harvard University, Cambridge MA, USA; Max Planck Tandem Group in Evolutionary Genomics of Specialized Metabolism, Universidad Nacional de Colombia, Bogotá, Colombia; Department of Genetics, University of Georgia, Athens GA, USA; Section of Plant Breeding and Genetics, Cornell University, Ithaca, NY USA 14853; Department of Biology, Boston College, Boston MA, USA

**Keywords:** self-incompatibility, mating system, *Phlox*, Polemoniaceae, QTL mapping, bulk segregant analysis, *S*-locus, sporophytic self-incompatibility (SSI)

## Abstract

Flowering plants display extensive variation in selfing rate, a trait with significant ecological and evolutionary consequences. Many species use genetic mechanisms to recognize and reject self-pollen (termed self-incompatibility or SI), and the loss of SI is one of the most common evolutionary transitions among flowering plants. Despite the ubiquity of transitions to self-compatibility (SC), little is known about the genetic architecture through which SC evolves. Specifically, it is important to determine if SC has a polygenic or simple genetic basis and if variation in compatibility localizes to the genomic locus causing self-pollen recognition (the “*S*-locus”). *Phlox drummondii* (Polemoniaceae) has been a model system for exploring mating system evolution and expresses extensive range-wide variation in the SI response. Here we investigate the genetic architecture of SC variants segregating within this otherwise SI species. Using multiple independent crosses, we uncover numerous QTLs associated with intraspecific variation in SI, consistent with a polygenic genetic architecture. While some QTLs overlap across mapping experiments, other QTLs are unique, suggesting that multiple genetic routes to SC exist. Through these crossing experiments, we demonstrate that *P. drummondii* has a sporophytic SI system, suggesting that an independent evolution of SI occurred in the lineage containing *Phlox*. We map this novel *S*-locus and find that the genomic region containing the *S*-locus is associated with intraspecific variation in SI in one of the three mapping populations. Although further work is necessary to clarify the conditions under which quantitative variation in SI represents a transitional pathway to complete SC, our study reveals the underlying genetic architecture upon which selection could act to drive this frequent and evolutionarily significant transition.

## 1. Introduction

The rate of self-fertilization within a species has wide-reaching impacts on evolutionary change. Through its control on effective population size, effective recombination, and population connectivity, a species’ mating system affects the rate and mode of evolution across the entire genome (Holsinger 2000; Igic *et al*. 2008; Wright *et al*. 2013). Most flowering plants (and even some animals) are co-sexual which provides the potential for self-fertilization; however, many species have evolved morphological, developmental, or genetic mechanisms to limit self-fertilization in response to the deleterious consequences of inbreeding (Barrett 2002; Porcher and Lande 2005; Charlesworth 2006; Anderson *et al*. 2010; Noël *et al*. 2017). The most widely employed strategy enforcing outcrossing in flowering plants is genetic self-incompatibility (SI), which involves the rejection of self-pollen deposited at the pistil based on its genetic identity (Silva and Goring 2001). Self-pollen recognition is controlled by a multi-allelic locus (the so-called “*S*-locus”) containing genes responsible for distinguishing self- vs. non-self-pollen (de Nettancourt 2010). Self-incompatibility has independently evolved many times throughout flowering plant evolution (Charlesworth *et al*. 2005; Allen and Hiscock 2008; Igic *et al*. 2008). Of the few well-characterized systems (e.g., Brassicaceae, Papaveraceae, and Solanaceae), each possesses an *S*-locus containing unrelated gene types, revealing that the common phenotype of self-incompatibility has diverse underlying molecular mechanisms across independent evolutions of SI (Nasrallah *et al*. 1985; Anderson *et al*. 1986; Foote *et al*. 1994; Zhang *et al*. 2024).

Despite the costs associated with selfing, the evolution of predominant self-fertilization from within a self-incompatible, outcrossing lineage is one of the most common evolutionary transitions in angiosperms (Stebbins 1974). The apparent ubiquity of such transitions may be explained by the broad and diverse scenarios under which selection favors self-fertilization in the short term. Mutations conferring selfing experience a genetic transmission advantage because selfing individuals act as both pollen and ovule donor for their selfed offspring while maintaining the ability to serve as an outcross pollen donor (Lloyd 1979; Stone *et al*. 2014). Furthermore, self-compatible individuals benefit from reproductive assurance when mating opportunities are rare (Baker 1955; Porcher and Lande 2005). A significant body of theoretical and empirical work has established ecological and geographic correlates of self-compatibility (SC) across diverse plant taxa including an association with small population size and peripheral or isolated habitats (Baker 1967; Herlihy and Eckert 2005; Busch 2005; Pannell *et al*. 2015; Grossenbacher *et al*. 2017; Koski *et al*. 2019). These patterns elucidate the selective pressures driving the evolution of self-fertilization and clarify why self-compatibility may be so common. However, establishing generalizations regarding the genetic basis of transitions to self-compatibility has been more protracted, in part because the genetic basis of self-incompatibility is so divergent across angiosperms, rendering candidate gene-based approaches inviable for most plant families (Zhang *et al*. 2024). To understand how self-compatibility evolves, we need to identify both the genetic basis of self-incompatibility across diverse plant groups and the genetic changes that facilitate its breakdown.

The evolution of self-compatibility in diploid systems can occur through two genetic routes: either through disruptive mutations at the *S*-locus or at unlinked genes that modify *S*-locus gene expression or encode downstream components of the pollen rejection pathway (Nettancourt 1977). In *Arabidopsis thaliana*, *Capsella rubella*, *C. orientalis*, and selfing-lineages of Siberian *A. lyrata*, mutations at the *S*-locus have caused the loss of self-incompatibility (Nasrallah *et al*. 2002; Guo *et al*. 2009; Bachmann *et al*. 2019; Kolesnikova *et al*. 2023). Analyses in the tomato clade (*Solanum* section *Lycopersicum*), attribute self-compatibility to variation at the *S*-locus (Markova *et al*. 2017; Broz *et al*. 2021). Similarly, in *Citrus, Petunia,* and *Prunus* mutations within the *S*-locus result in self-compatibility (Tsukamoto *et al*. 2003b; Liang *et al*. 2020; Li *et al*. 2020). These numerous case-studies span diverse taxa and types of self-incompatibility systems and together suggest that mutations at the *S*-locus may be a common genetic route to self-compatibility. However, unlinked modifier loci have also been implicated (McClure *et al*. 1999; Nasrallah *et al*. 2004; Liu *et al*. 2007). For example, in North American *A. lyrata*, when inherited with a specific *S*-allele, a dominant-acting unlinked modifier can disrupt the self-recognition response (Li *et al*. 2023). In *Citrus, Petunia,* and *Prunus,* modifier loci are similarly involved in modulating expression of *S*-locus genes (Tsukamoto *et al*. 2003a; Ono *et al*. 2018; Hu *et al*. 2021). Distinguishing between these two possible genetic architectures is critical for understanding how selection may act during mating system evolution. Several theoretical models describing when and how mating system evolution occurs demonstrate that the genetic architecture of self-compatibility affects evolutionary outcomes (Lande and Schemske 1985; Charlesworth and Charlesworth 1990; Latta and Ritland 1993). For example, under certain evolutionary conditions, intermediate levels of self-incompatibility can stably evolve within a species when self-compatibility is caused by a modifier loci unlinked to the *S*-locus (Porcher and Lande 2005).

Although self-incompatibility is often treated as a categorical phenotype, it is increasingly appreciated that variation in self-pollen rejection is quantitative (Stephenson 2000; Whitehead *et al*. 2018). Heritable variation in the self-incompatibility response (sometimes termed pseudo-self-compatibility or partial self-incompatibility) has been shown within several plant species including *Campanula rapunculoides*, *Leavenworthia alabamica*, *Solanum habrochaites*, and *Witheringia solanacea* (Good-Avila and Stephenson 2002; Stone *et al*. 2006; Baldwin and Schoen 2017; Broz *et al*. 2017). Mapping the genetic basis of such intraspecific variation has advantages over mapping fixed differences between completely SI and completely SC species. Once a mutation disrupts the SI response, other SI genes become unnecessary and relaxed purifying selection can permit numerous loss-of-function mutations to accumulate within a completely SC species. Therefore, identification of the genetic basis of divergent compatibility phenotypes between species may not uncover the original mutation(s) upon which selection acted. Additionally, in one of the most extensively studied examples of a mating system transition, the highly-selfing *A. thaliana,* genomic evidence supports at least three independent origins of self-compatibility, indicating that several SC mutations were segregating during the transition (Shimizu and Tsuchimatsu 2015). If multiple SC mutations segregate within a species, not all necessarily go on to become fixed differences between species. Intraspecific variation in the self-rejection response can help identify the genetic material through which selection acts to drive mating system evolution.

The wildflower, *Phlox drummondii* (Polemoniaceae), is an ideal system to investigate the genetic basis of within-species variation in self-incompatibility. *Phlox* has been used as a model for understanding the ecological and evolutionary implications of mating system variation (Levin 1978, 1989, 1993, 1996; Clay and Levin 1989; Plitmann and Levin 1990; Bixby and Levin 1996). *Phlox drummondii* expresses quantitative variation in the self-pollen rejection response and this trait is highly heritable (h^2^ = 0.47-0.90; (Bixby and Levin 1996; Roda and Hopkins 2019). Very little is known about the genetic or molecular basis of the self-pollen rejection response in *P. drummondii* (or any other member of Polemoniaceae) rendering candidate gene-based approaches infeasible. Alternatively, classic quantitative genetic studies can untangle the type of genetic mechanism conferring self-recognition and the genetic basis of variation in self-pollen rejection.

Self-incompatibility systems in flowering plants can be classified into two categories: gametophytic self-incompatibility (GSI) and sporophytic self-incompatibility (SSI). These phenotypic categories are used to organize the diversity of SI systems based on the life-stage at which the pollen genotype is recognized (Nettancourt 1977). In GSI, the pistil recognizes the haploid *S*-genotype of the pollen such that each pollen grain has a single *S*-allele specificity. In SSI, the pistil expressed gene recognizes the diploid *S*-genotype of the pollen-producing parent such that each pollen grain may carry the biallelic *S*-specificity of the anther from which it developed. Early work in *P. drummondii* suggested that self-incompatibility is controlled by a gametophytic type system (Levin 1993). Subsequent work in a related Polemoniaceae species, *Linanthus parviflorus*, pointed to a sporophytic type system (Goodwillie 1997). There are no other documented examples of both GSI and SSI existing within a single plant family, suggesting that mating system evolution is either exceptionally labile within the Polemoniaceae or that the interpretation of previous crossing data was confounded. Controlled crosses have classically been used to differentiate GSI and SSI based on patterns of cross-compatibility among full-siblings (Thompson and Taylor 1966; Nettancourt 1977). Segregating variation causing a loss of self-incompatibility within a species can disrupt predictable patterns of incompatibility within a crossing family and make it challenging to accurately distinguish GSI from SSI. Therefore, determining the type of self-incompatibility system active in *Phlox* while simultaneously investigating the genetic architecture of variation in the self-incompatibility response is necessary.

Here, we examine the genetic basis of intraspecific variation in self-incompatibility within the wildflower species, *P. drummondii*. We describe quantitative variation in self-incompatibility across the species’ range and use a traditional quantitative trait locus mapping experiment paired with multiple bulk segregant analyses to identify quantitative trait loci (QTLs) underlying this intraspecific variation. We perform genetic mapping across several independent crosses to explore whether multiple mutations causing a loss of self-incompatibility are segregating within *P. drummondii* and to characterize their genetic architecture. By combining a diallel crossing experiment with bulk segregant mapping, we also identify the genomic region containing the *S*-locus for the first time in the phlox family and definitively demonstrate sporophytic control of self-pollen recognition in *Phlox*. Finally, we quantify gene expression patterns in reproductive organs to generate a list of candidate genes underlying our QTLs. Collectively, our work reveals the genetic architecture of mutations contributing to the evolution of self-compatibility and allows us to determine if the loss of self-incompatibility involves mutations at the *S*-locus and/or at unlinked genetic modifiers in this system.

## 2. Methods

### Study species

*Phlox drummondii* (Polemoniaceae) is an annual wildflower native to central Texas, USA. This species has an active self-incompatibility system, and there is natural quantitative variation in the expression of self-incompatibility as measured by the number of seeds that set following manual self-pollination (Roda and Hopkins 2019). This variation is heritable and responds to artificial selection (Bixby and Levin 1996). While rates of autonomous selfing in both greenhouse and field conditions are low (<2%), geitonogamous visitation by the primary pollinator, *Battus philenor*, is common and results in frequent transfer of self-pollen between flowers (Burgin *et al*. 2023).

### Plant collection and care

In May of 2018 and 2019, we collected seeds from naturally occurring *P. drummondii* individuals throughout the species range in Texas, USA. Plants were grown under standardized conditions at the Arnold Arboretum of Harvard University during 2020 and 2021. Specifically, we incubated seeds with 500 ppm gibberellic acid in water for two days and vernalized planted seeds at 4^0^C for ten days. After germination, plants were transferred to 4-inch pots with Pro-Mix HP Mycorrhizae potting media and maintained in growth chambers. Plants were fully randomized to pot location and trays were rotated regularly. Growth chambers were kept at a daytime temperature of 23^0^C and a nighttime temperature of 18^0^C, with 16 hours of supplemented light. All mapping populations used in our analyses were produced from crosses of these plants and grown in growth chambers under the same standardized conditions.

### Quantifying variation in self-incompatibility across the species range

To document natural variation in self-incompatibility, we expanded on the dataset presented by Roda and Hopkins (2019) using consistent methods of controlled pollinations on greenhouse-grown plants. Corollas with anthers attached were removed one to three days prior to anther dehiscence/floral opening. Three days after anther removal, we used forceps to deposit pollen on fully developed stigmas. Controlled crosses were covered in fine mesh bags to prevent seed loss following explosive fruit dehiscence. Previous work demonstrated that the number of self-pollen grains germinating at the stigmatic surface predicts the number of seeds that will form following self-pollination (Roda and Hopkins 2019). Self-seed set is therefore a reliable metric for self-rejection occurring at the pollen-pistil interface in this system. After a minimum of two weeks following pollination, we removed mesh bags and counted the total number of seeds produced. *Phlox drummondii* flowers contain three ovules, and a single cross will therefore produce between zero and three seeds (though flowers occasionally produce four ovules and therefore up to four seeds in a cross).

To explore quantitative variation in this trait, we crossed eight flowers per plant with self-pollen, which resulted in zero to 25 seeds per plant in our study. We counted the number of seeds produced following pollination with self-pollen as a measure of the strength of self-incompatibility (i.e., individuals producing more seeds following self-pollination are more self-compatible). Individuals originally presented in Roda and Hopkins (2019) were phenotyped using a range of number of flowers per plant; to maximize consistency with our current dataset, we only included individuals phenotyped using eight flowers per plant. To control for variation in plant fertility and health unrelated to self-incompatibility, we used the same method to cross an additional eight flowers with pollen derived from an unrelated *P. drummondii* plant chosen at random (outcross pollinations). Any individual that produced less than two-thirds of full seed set (e.g., fewer than 16 seeds) following outcross pollination was removed from the experiment.

### Genetic mapping

We used a series of quantitative trait locus (QTL) mapping experiments to identify regions of the genome associated with intraspecific variation in self-incompatibility as well as the region of the genome corresponding to the self-pollen recognition *S*-locus. These included one experiment using traditional QTL mapping analyses and four bulk segregant analysis (BSA) experiments as described below. Both approaches highlight the genomic position of QTLs and provide unique advantages. We used a traditional QTL mapping approach which enables us to determine the magnitude and direction of genotypic effects at each QTL. While BSA cannot provide information about the phenotypic effects of QTLs, it has long been used to efficiently gain genome-wide information at low-cost. Therefore, to test whether multiple mutations conferring self-compatibility are segregating within *P. drummondii*, we leveraged this feature of BSA to map variation in SI in multiple independent crosses.

#### (a) QTL mapping of variation in self-incompatibility

For our traditional QTL mapping population, parental plants included one fully self-incompatible individual collected as seed from a naturally occurring population, Population 633 (30.0505, -97.2378) and one highly self-compatible individual collected as seed from a different naturally occurring population, Population 608 (30.1334, -97.101). These plants were crossed to generate a single F1 individual. Immature floral buds of self-incompatible plants are self-compatible; we leveraged this developmental variation to generate F2 individuals by repeatedly self-fertilizing our F1 individual. For clarity, we will refer to this mapping population as “Family A.” F2 individuals were germinated and grown as described above. We performed eight controlled self-pollinations and eight controlled outcross pollinations on each F2 individual. Plants setting fewer than 15 outcross seeds were removed from the experiment, resulting in a final mapping population of 169 F2 individuals. We used total self-seed set as a quantitative measure of self-incompatibility and performed QTL mapping for this trait.

#### (b) Bulk segregant analysis of variation in self-incompatibility

To generate our BSA mapping populations (called “Family B” and “Family C” for clarity), we randomly selected two plants grown from seed collected in two geographically distinct, naturally occurring populations: individual B from Population 664 (30.116, -97.365) and individual C from Population 667 (29.5288, -96.4412). These individuals were self-fertilized to generate 93 and 220 S2 individuals in families B and C respectively. We performed six controlled pollinations per individual using self-pollen and monitored for fruit set. Plants for each family were divided into two incompatibility groups: self-incompatible and self-compatible. Plants setting zero self-fruits were categorized as self-incompatible. The threshold for self-compatibility was determined based on each family’s fruit set distribution; plants setting at least one self-fruit in Family B considered self-compatible and plants setting at least three self-fruits in Family C were categorized as self-compatible (Figure 1).

**Figure 1.**
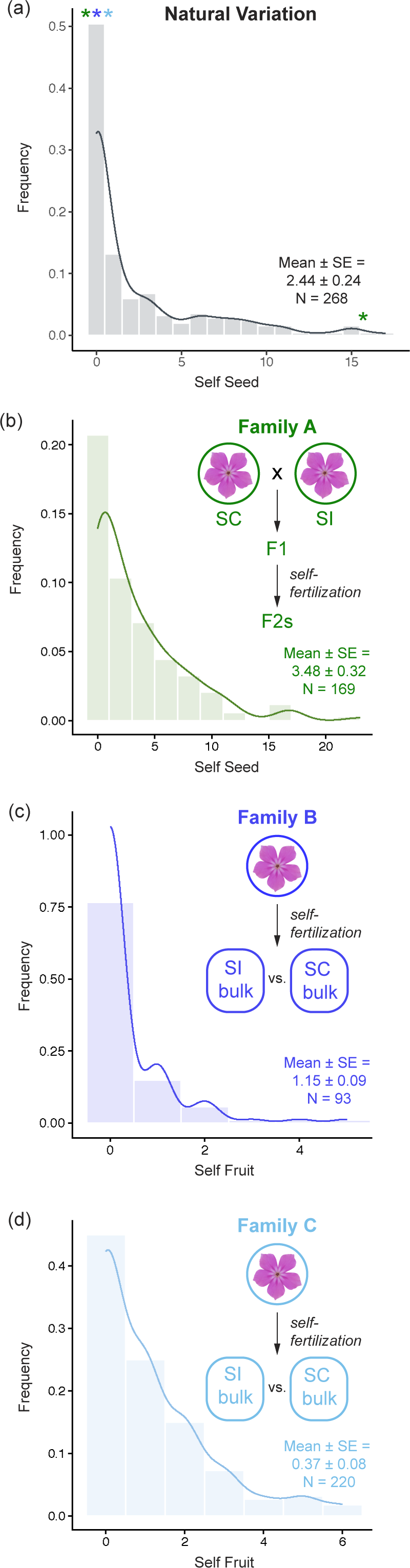
Variation in self-incompatibility within *Phlox drummondii* as measured by seed or fruit set following self-pollination. Lines depict the density curves for each distribution. **(a)** Individuals grown from field-collected seed have quantitative variation in the self-incompatibility response under greenhouse conditions. Asterisks indicate the phenotype value for plants selected to create Families A (green), B (dark blue), and C (light blue). **(b-d)** Graphical summaries of cross design are shown above the phenotypic distributions for mapping individuals. F2s individuals are plotted for Family A and S2 individuals for Families B and C.

#### (c) Bulk segregant analysis of the *S*-locus in P. drummondii

Previous work in *P. drummondii* suggests that recognition of self-pollen is controlled by a single genetic locus (the “*S*-locus”). However, the genetic identity of the *S*-locus in *Phlox* or any other member of Polemoniaceae is unknown. To test whether variation at the *S*-locus is involved in intraspecific variation in self-incompatibility, we used a bulk-segregant mapping approach to identify the genomic region containing the *S*-locus in *Phlox*. We performed *S*-locus mapping using only the self-incompatible individuals from the previously described families B and C. Experiments were performed separately for each family.

We used a diallel crossing design to group individuals by their genotype at the *S*-locus. Because S2 offspring were generated through self-fertilization, individuals in each family are one of three possible genotypes at the *S*-locus: heterozygote or alternative homozygotes. Crosses between individuals with the same homozygous *S*-locus genotype are recognized as self and result in no fruit set in reciprocal crosses. Simultaneously, crosses between individuals with different homozygous *S*-locus genotypes are recognized as non-self and fruits will develop in reciprocal crosses. Heterozygous individuals are expected to have variable patterns of incompatibility depending on if the *S*-locus system recognizes the haploid genotype of each pollen grain (gametophytic incompatibility) or the diploid genotype of the pollen producing plant (sporophytic incompatibility). In sporophytic systems, dominance among *S*-alleles can further complicate expectations of incompatibility for heterozygous individuals. We began by conducting a full diallel cross with 10 S2 individuals from within each mapping family. After identifying groups of plants with similar patterns of cross-compatibility (and therefore *S*-locus genotype), we subsequently categorized each remaining individual by performing reciprocal crosses with one plant from each crossing/genotype group.

### DNA extraction and genotyping

For all individuals in the mapping populations, we extracted DNA from fresh leaf tissue using EZNA Plant kit buffers (Omega Bio-Tek) followed by a modified chloroform extraction and two cold 96% ethanol washes as previously described (Goulet-Scott *et al*. 2021). For the bulk segregant populations, we combined an equal quantity of genomic DNA from individuals within a group for bulk sequencing. To map variation in self-incompatibility, we bulked individuals within a family depending on their self-incompatibility phenotype. To map the *S*-locus, we bulked individuals within a family that were members of the same homozygous *S*-locus genotype and did not include heterozygotes for this analysis.

We constructed libraries for each individual of the QTL mapping population Family A and for each bulk in Families B and C using a double-digest restriction site-associated (ddRAD-seq) protocol. Libraries were then pooled, size-selected for fragments with length 300-500bp using a Pippin Prep (Sage Biosciences) and cleaned with a Monarch PCR and DNA cleanup kit (New England Biolabs). Samples were paired end sequenced (2x150bp) on an Illumina NovaSeq 6000 at the Bauer Core of Harvard University.

Raw reads were demultiplexed and filtered as described in Goulet-Scott at al., 2021. Reads were aligned using BWA-MEM to a complete assembly of the *P. drummondii* genome. Individuals and bulks were genotyped using the *gstacks* and *populations* tools in STACKS v2.64 to create individual or joint cohort-level gVCFs. Average read depth per sample was calculated using VCFtools. To control for overall sample quality, any individual or bulk with a mean read depth below 10 was removed from the analyses. To prevent the inclusion of low-quality genotypes in our study, sites were filtered using VCFtools. Specifically, sites were filtered to include only biallelic SNPs with no more than 50% missingness across individuals. Genotypes were further filtered to include only those with genotype quality greater than or equal to 15.

### RNA sequencing

We collected five unique plant tissues for RNA sequencing: stigma and style (pistil with the ovary removed), pollen, anther, leaf, and stem. Our RNA-seq experiment totaled eighteen samples including six biological replicates for stigma and style samples and three biological replicates for all other tissues (Fig. S4). Each stigma and style sample included tissue pooled from eighteen flowers of the same plant. To prevent self-pollen contamination and developmentally standardize flower age, we removed petals (with non-dehisced, attached anthers) from floral buds predicted to open in three days and collected tissue following those three days. All tissue was flash-frozen in liquid nitrogen to prevent RNA degradation.

We extracted total RNA using the Spectrum Plant Total RNA Kit (Sigma-Aldrich) with an additional DNA removal step using the Qiagen on-column DNAse kit. We used a NanoDrop 8000 spectrophotometer (Thermo Fisher Scientific) and a 2200 TapeStation (Agilent Technologies) to assess RNA concentration and quality. The Bauer Core Facility at Harvard University prepared libraries using a KAPA stranded RNA HyperPrep Kit (Roche Sequencing) with Illumina adapters. Samples were paired end sequenced (2x150bp) on an Illumina NovaSeq 6000 at the Bauer Core of Harvard University (NCBI SRA SUB15344310). Following adapter trimming using Trim Galore! V0.6.2, reads were aligned to a complete assembly of the *P. drummondii* genome (NCBI BioProject: PRJNA1219593, unpublished) using a 2-pass alignment with STAR v2.7.11b (Dobin *et al*. 2013; Kreuger 2019). Abundance calls were generated using RSEM v1.3.0 and used as the input for downstream expression analyses (Li and Dewey 2011).

### Statistical Analyses

#### (a) Linkage map construction and QTL mapping of variation in self-incompatibility

We used Lep-Map3 suite v0.5 to construct a linkage map for *P. drummondii* using the QTL mapping population described above (Rastas 2017). Lep-Map3 is designed for F2 populations produced by crossing two unique F1 individuals. However, F2 individuals in our study were produced via self-fertilization of a single F1. To account for this design, we constructed our pedigree file to include a “dummy Male” and “dummy Female” individual whose parents were specified as the two parents in the original cross. We removed the actual F1 from the analysis and treated the “dummy Male” and “dummy Female” as parents to the F2 individuals. To generate parental genotype markers, we passed the pedigree and VCF files to the *ParentCall2* module. We then used the *Filtering2* module to filter markers under the following parameters: removeNonInformative = 1, dataTolerance = 0.0001, MAFLimit = 6, missingLimit = 0.5. Markers were then assigned to linkage groups using the *SeparateChromosomes2* module under the following parameters: lodLimit = 50, theta = 0.0013, sizeLimit = 50, grandparentPhase = 1. We then used *JoinSingles2All* to incorporate previously unassigned markers into the main linkage groups under the same lodLimit and theta parameters as before. Finally, we used the *OrderMarkers2* module to phase markers within each linkage group with outputPhasedData = 1.

QTL mapping of variation in the self-incompatibility response was performed in R/qtl using total self-seed set as a quantitative trait. We calculated conditional genotype probabilities using the *calc.genoprob* function with a fixed step size of 2cM, an assumed genotyping error rate of 0.05, and the Kosambi mapping function. The *scanone* function was used with the HK algorithm under a nonparametric model, and we established significance thresholds using permutations (n=1000). We consider *p* < 0.05 as significant. For significant QTLs, the percentage of variation explained (PVE) was calculated using the *fitqtl* function with the HK algorithm.

#### (b) Bulk segregant analyses

All analyses were implemented using the QTLseqR package in R and conducted separately for Families B and C. Genomic markers were selected after aligning reads to the genome and filtering as described under the methods section “DNA extraction and genotyping.” Allele frequencies are expected to vary between bulks for markers linked to loci affecting self-incompatibility while unlinked markers will have an allele frequency difference near zero. Following this logic, we used 50 Mb sliding windows across the genome to compare allele frequencies between SI and SC bulks. We used 100 Mb sliding windows in the *S*-locus genotype comparisons because we expected a single peak consistent with a single locus causing self-pollen recognition specifically. To account for sampling effects of read depth and bulk size, we used a nonparametric test of allele frequency divergence between bulks. More specifically, we calculated a modified G-statistic smoothed by a weighted average procedure to produce a statistic for each sliding window (G’) (Magwene *et al*. 2011). To establish significance thresholds, we adjusted p-values to correct for multiple testing at a false discovery rate of 0.05.

#### (c) Identifying candidate genes using gene expression

We identified candidate genes underlying the two most significant QTLs from each of our mapping populations. To establish genomic intervals for each QTL in Family A, we used the base pair positions of markers at the start and end of 1.5 LOD intervals for each QTL. In Families B and C, we used the genomic interval for which all SNPs pass the false-discovery threshold of q < 0.05. We then identified genes within these regions based on the gene annotation for the *P. drummondii* genome. Because the SI response occurs only in the stigma/style, we reasoned that genes involved in the SI pathway are likely restricted in expression to stigma/style tissue. To identify genes following this pattern, we normalized read counts across samples using the TMM method and characterized differentially expressed genes pairwise between stigma/style samples and each other tissue type independently in edgeR (Robinson *et al*. 2010). We used the quasi-likelihood *F*-test to identify differentially expressed genes using the glmQLFTest and adjusted p-values with the Benjamini-Hochberg Procedure. These gene lists were filtered to include only genes with greater than 5-fold-change and a significance threshold of p < 0.01. We considered a gene to be specifically expressed in stigma/style tissue if it met these criteria in all pairwise tissue comparisons. We conducted a parallel analysis for anther tissue and considered a gene to be specifically expressed in the anther if it met these criteria in pairwise tissue comparisons with stigma/style, leaf, and stem. Comparison with pollen was omitted from this analysis.

## 3. Results

### Phenotypic variation among field-collected plants

We find quantitative variation in self-incompatibility within *P. drummondii* through controlled pollinations on 268 individuals from 38 naturally occurring populations spanning the species range (Fig. 1A). While most individuals set zero self-seeds, approximately 40% of individuals set at least one self-seed. Some degree of self-compatibility is present in all sampled populations but one (Population 675: 29.4024, -97.774517, n=4, Fig. S1; Table S1).

### Genetic architecture underlying intraspecific variation in self-incompatibility

After filtering, Family A F2s were sequenced at an average read depth of ∼44x with 22,925 sites and a missingness of 12.3% per individual. Our final linkage map contains 6,992 markers in seven linkage groups (LGs) consistent with the expected chromosome number in *Phlox*. The total map length is 795.4cM with an average marker distance of 0.1cM (Table S2). To identify the genetic architecture of intraspecific variation in self-incompatibility, we scored 169 recombinant SIxSC F2 individuals for self-seed set. As observed among field-collected plants, we find quantitative variation in the number of seeds formed following self-pollination; self-seed set ranges from 0-23 seeds (out of 24 possible) with most individuals setting no self-seed (Fig. 1B). We mapped the genetic basis of this variation and identify a single QTL on LG3 (Q3A) passing genome-wide significance (Fig. 2A; Fig. S3). Two additional QTLs on LG1 (Q1A) and LG5 (Q5A) do not meet genome-wide significance but are significantly associated in an individual marker analysis (Q1: F-value=4.917, p<0.008; Q2: F-value=3.865, p<0.023; Fig. S3). No significant interactions between these QTLs were identified. The model including all three QTLs had a total LOD score of 7.83 and collectively explains 19.21% of phenotypic variance in self-seed set (Table 1).

**Figure 2.**
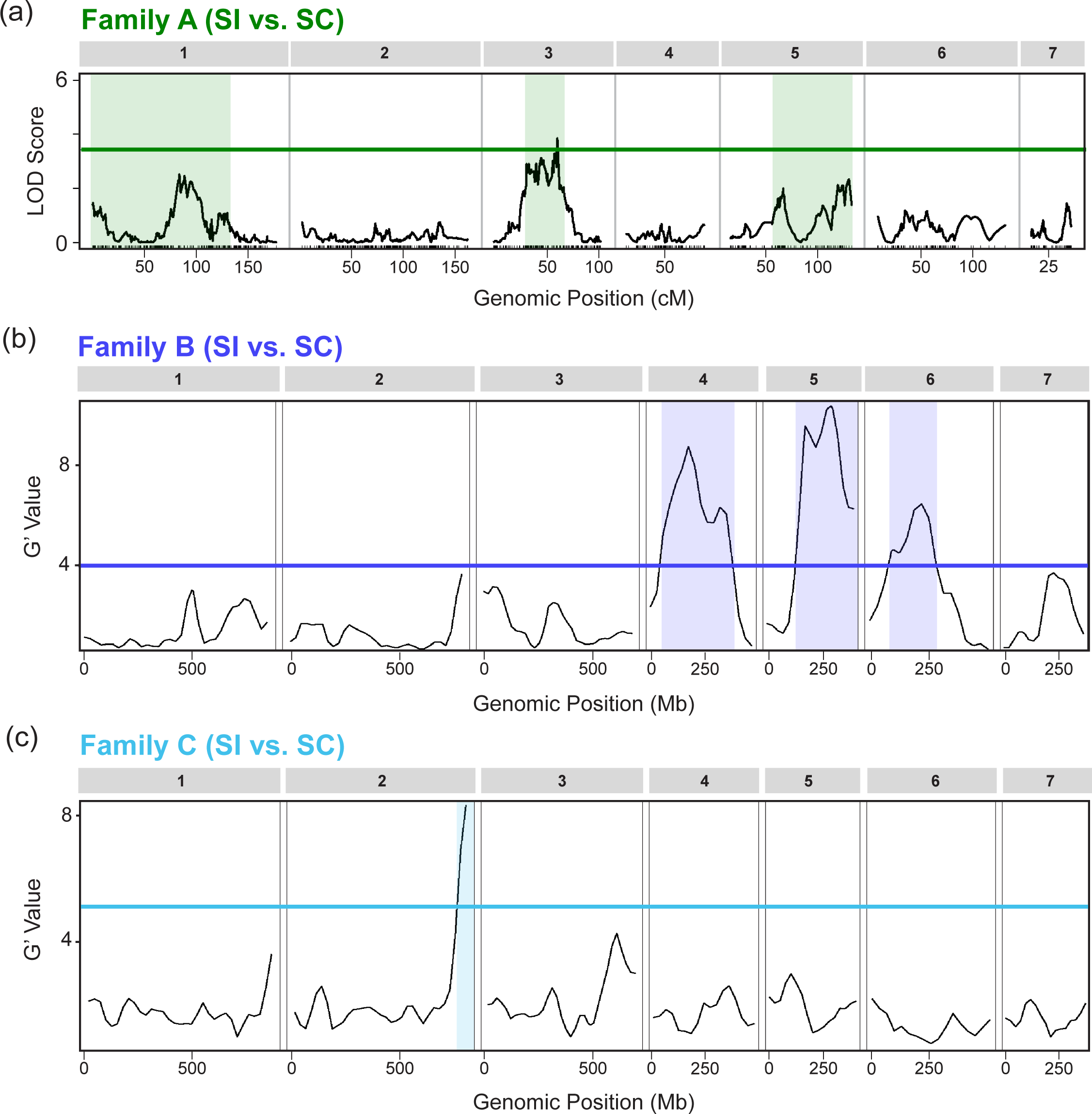
The genetic architecture of quantitative variation in the self-incompatibility response. **(a)** Strength of association, as indicated by LOD scores, between genotype and self-seed set in Family A F2 individuals across the seven chromosomes of *P. drummondii.* Green line indicates a genome-wide significance cutoff of 0.05 based on 1000 permutations. **(b)** Association between deviation in allele frequence and self-incompatibility, as indicated by a G’ statistic, across the seven chromosomes in *P. drummondii* in Family B and **(c)** Family C. Blue lines indicate a q-value significance cutoff of 0.05 for each mapping population

**Table 1.** Summary statistics for Family A QTLs.

| QTL | Linkage Group | 1.5 LOD int (cM) | Peak Position (cM) | Peak LOD | PVE (%) | Additive Effect (SE) | Dominance Effect (SE) |
| --- | --- | --- | --- | --- | --- | --- | --- |
| Q3A | 3 | 37.21-69.62 | 67.4 | 3.84 | 8.13 | -1.27 (0.40) | 1.54 (0.59) |
| Q1A | 1 | 22.41-152.6 | 106.2 | 2.16 | 5.04 | 0.72 (0.41) | -1.53 (0.59) |
| Q5A | 5 | 89.11-165.97 | 163.26 | 1.71 | 4.07 | -0.27 (0.40) | 1.59 (0.60) |

Abbreviations: LOD, Logarithm of the Odds; int, interval; PVE, Percent Variance Explained

We performed mapping experiments in two additional, independent crosses using a bulk-segregant analysis approach (Fig. 1C&D; Table S3). In Family B, we find three QTLs significantly associated with variation in self-incompatibility on LGs 4, 5, and 6 (Q4B, Q5B, and Q6B respectively). In Family C, we find a single QTL (Q2C) that passes genome-wide significance located on LG2 (Fig. 2C; Table 2).

**Table 2.** Summary statistics for Family B and C bulk segregant associations.

| Comparison | Family | QTL | Linkage Group | Start | End | Max G' | Mean G' | Mean q-value |
| --- | --- | --- | --- | --- | --- | --- | --- | --- |
| SI Variation | B | Q5B | 5 | 151,078,507 | 405,593,763 | 12.896 | 8.746 | 0.004 |
| SI Variation | B | Q4B | 4 | 59,701,670 | 362,380,151 | 10.882 | 8.406 | 0.004 |
| SI Variation | B | Q6B | 6 | 188,317,564 | 281,801,628 | 8.041 | 7.648 | 0.005 |
| SI Variation | C | Q2C | 2 | 781,731,802 | 799,378,250 | 8.322 | 7.707 | 0.001 |
| *S*-Genotype<br>(*S1S1* vs. *S2S2*) | B | *S*-locus | 2 | 732,745,439 | 799,378,250 | 15.166 | 13.699 | 0.024 |
| *S*-Genotype<br>(*S3S3* vs. *S4S4*) | C | *S*-locus | 2 | 726,864,093 | 799,378,250 | 13.329 | 11.616 | 0.015 |

Collectively, these mapping experiments demonstrate that the genetic architecture of variation in self-incompatibility is polygenic. We find overlapping QTLs in independent crosses when using individual marker association analyses, yet most QTLs appear in only one crossing experiment, demonstrating that multiple genetic routes to self-compatibility exist within *P. drummondii* (Fig. 2).

### Identifying the *S*-locus in Phlox

After excluding self-compatible individuals, we recover three cross-compatibility groups in both Family B and C, consistent with three possible *S*-locus genotypes segregating in each family (Fig. 3A&B). Compatibility groups in both families display reciprocal differences in incompatibility; the inferred heterozygous group is compatible as a pollen donor but not as a recipient with one or more of the homozygous classes (Fig. 3A&B). These patterns of cross-compatibility cannot be explained by gametophytic control of self-pollen recognition. However, our crossing results for both families are consistent with sporophytic self-incompatibility. We mapped the genomic position of the *Phlox S*-locus by comparing allele frequencies between the inferred alternative bulked homozygous genotypes in each family (Table S3). In both families, we find a single peak on LG2 that significantly associates with inferred *S*-locus genotype (Fig. 3C&D). This region overlaps with the QTL associated with variation in self-incompatibility in Family C, suggesting that mutations at the *S*-locus contribute to self-compatibility but are not the only genetic route to self-compatibility within this species.

**Figure 3.**
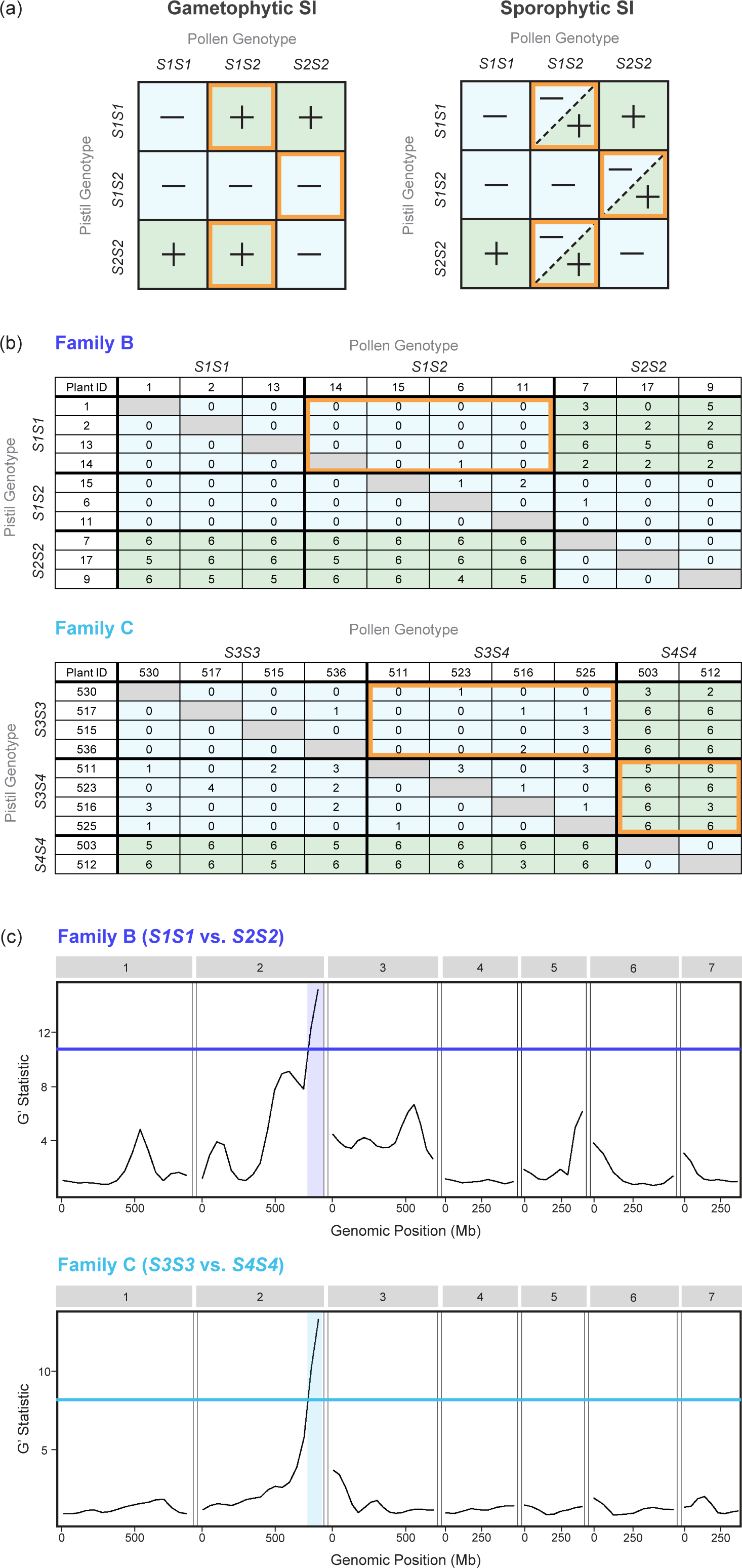
Two independent crosses reveal that self-pollen recognition in *Phlox* is sporophytic with dominance and controlled by a single genomic locus (the *S*-locus) located on LG2. **(a)** Schematic representations of the expected crossing results for gametophytic self-incompatibility (GSI) and sporophytic self-incompatibility (SSI). Compatible crosses are indicated with a plus sign and shaded light green while incompatible crosses are indicated with a minus sign and shaded light blue. In GSI, heterozygous pollen should be compatible with both homozygous pistil genotypes. In SSI, heterozygous pollen may or may not be compatible with either homozygous pistil genotype, depending on the dominance hierarchy among *S*-haplotypes. Orange boxes refer to crossing results that can distinguish SSI from GSI. **(b)** Crossing results for the completely SI subsets of Families A and B. In Family A, heterozygous pollen is incompatible with *S1S1* homozygous pistils (noted by an orange box), a pattern consistent with SSI with dominance (*S2* is dominant to *S1* in pollen). In Family B, heterozygous pollen is incompatible with *S3S3* homozygous pistils and *S4S4* homozygous pollen is compatible with heterozygous pistils (noted by orange boxes). This cross-compatibility pattern is also consistent with SSI with dominance (*S3* is dominant to *S4* in pollen and pistil). **(c)** G’ statistic indicating association between deviation in allele frequency and *S*-locus genotype class across the seven linkage groups in *P. drummondii*. Blue lines indicate a q-value significance cutoff of 0.05.

### Candidate genes

Our PCA of normalized reads demonstrates clear clustering by tissue type, with PC1 and PC2 together explaining 34.03% and 15.75% of the overall variance, respectively (Fig. S4). There are 224 total genes with stigma/style specific expression. We find eight genes with stigma/style specific expression within the 1.5 LOD interval for Q3A out of 1,779 total genes annotated to this region in the *P. drummondii* reference genome (Table S4). One candidate gene, PD.55820, occurs within 1Mb of the peak marker. The closest *Arabidopsis thaliana* homolog to PD.55820 is implicated in the formation of reactive oxygen species, which are known to underly self-incompatibility responses in other sporophytic SI mechanisms (Zhang *et al*. 2021; Goring *et al*. 2023). Within the 1.5 LOD interval for Q1A, we find eleven genes with stigma/style specific expression out of 1,228 total genes annotated to this region, and nine genes out of 1,635 total annotated to Q5A. The three QTLs passing significance in Family B collectively contain nine stigma/style specific genes with three overlapping with those identified in Q5A (Fig. 2; Table S4&S5). The genomic region associated with *S*-locus genotype contains 727 total genes with eight restricted in expression to the stigma/style. This region partially overlaps with Q2C identified in the Family C map for variation in self-incompatibility which contains seven genes with restricted expression (Fig. 2&3; Table S4&S5). Notably, the closest *A. thaliana* homolog for one such stigma/style specific gene, PD.52568, is predicted to encode a receptor-like tyrosine kinase similar to the *S*-locus in Brassica, a very distantly related genus also expressing sporophytic self-incompatibility (Nasrallah *et al*. 1985; Stein *et al*. 1996; Igic *et al*. 2008). There are 845 total genes with anther specific expression and 145 of these are annotated within an identified QTL (Table S6&S7).

## 4. Discussion

Mating system variation has long been a central focus of the botanical literature, revealing the fundamental importance of this trait to ecological and evolutionary dynamics (Darwin 1876; Fryxell 1957; Barrett 2003; Charlesworth 2003; Goldberg *et al*. 2010; de Vos *et al*. 2014). Historical descriptions organized plant mating systems qualitatively, using the discrete categories of complete SI or SC. In reality, many plant species vary quantitatively in their expression of SI due to both genetic and environmental factors (Stephenson 2000; Travers *et al*. 2004; Raduski *et al*. 2012; Steinecke *et al*. 2022; Ferrer *et al*. 2024). Our work characterizes intraspecific, quantitative variation in SI that is heritable, range-wide, and has a polygenic basis. We map the genomic position of a novel *S*-locus and reveal that intraspecific variation in SI is associated with genetic variation at the *S*-locus as well as at unlinked loci. Previous work investigating the genetic basis of mating system variation relied mostly on between species crosses (Markova *et al*. 2017; Bachmann *et al*. 2019; Liang *et al*. 2020). All quantitative genetic studies capture variation at a specific moment during a much longer evolutionary process. Interspecific studies reveal the culmination of evolutionary forces acting on traits over thousands to millions of years and can confound the initial causal mutations with subsequent degradation of the incompatibility pathway via relaxed purifying selection. Evolutionary transitions to complete SC clearly require heritable, individual-level variation in the SI response and understanding the genetic architecture on which selection acts to drive these transitions necessitates a within-species approach. We leverage such intraspecific variation to identify a complex genetic architecture of SC variants segregating within an otherwise SI species. We demonstrate that phenotypic variation in this trait is highly quantitative, polygenic, and find unique loci segregating in independent crosses. The finding of multiple independent loci is perhaps unsurprising given previous estimates that SC mutations can arise within SI species at a non-negligible rate of approximately 2x10^-7^ to 1x10^-5^ per pollen grain (Nettancourt 1977; Lewis 1979). Whether these mutations will be positively selected to drive a transition to complete SC, maintained at intermediate frequency, or ultimately purged depends on complex ecological and evolutionary factors including the strength of inbreeding depression, effective population size, and the dominance/recessivity pattern of any given mutation (Nagylaki 1976; Lloyd 1979; Porcher and Lande 2005). Our findings motivate future research in *P. drummondii* to quantify these ecological and evolutionary factors, and to compare the variation identified in the present study with the genetic basis of complete SC in the closely related species, *P. cuspidata* (Levin 1978).

Fully functional SI depends on the action of genes within the *S*-locus responsible for self-pollen recognition as well as elements in the signaling cascade leading to rejection. By breaking the ability to distinguish self- and non-self-pollen, mutations at the *S*-locus will cause SC, and it has been shown that many fully SC species indeed have non-functional S-loci (Nasrallah *et al*. 2002; Tang *et al*. 2007; Guo *et al*. 2009; Bachmann *et al*. 2019; Kolesnikova *et al*. 2023). However, mutations in various components of the rejection pathway can also lead to SC. Quantitative variation in the SI response (sometimes termed “pseudo-self-compatibility”) has previously been assumed to be caused by mutations outside of the *S*-locus (Levin 1996). Interestingly, we find that this prediction is true within *P. drummondii* in some cases, but that genetic variation at the *S*-locus can also contribute to pseudo-self-compatibility. This finding suggests that the presence of quantitative variation in the SI response in other species may not be predictably caused by mutations either inside or outside the *S*-locus. Future work in *P. drummondii* should focus on characterizing the specific molecular mechanism through which mutations outside of the *S*-locus cause a reduction in the SI response. For example, do these mutations modify expression of genes encoded within the *S*-locus or do they impact function of the downstream signaling pathway without affecting *S*-locus expression (Li *et al*. 2023)?

Given the diverse molecular mechanisms underlying the three relatively well-characterized SI types, it is difficult to confidently suggest candidate genes underlying variation in the SI response (Zhang *et al*. 2024). With this limitation in mind, we present a list of genes within identified QTLs that follow patterns of tissue expression consistent with a mechanism operating only in pistil tissue (e.g., rejection of self-pollen). These candidates are particularly relevant for identifying cis-acting mutations underlying variation in SI. We additionally present a list of anther specific genes within the identified QTLs that will be useful candidates if future studies reveal that variation in SI is mediated by proteins acting on the pollen side of the post-pollination process. However, our QTL regions are physically large and contain many genes with plausible effects on the SI response. Further fine-mapping and functional genetic studies will be necessary to substantiate a possible role for the candidate genes presented here.

The phlox family (Polemoniaceae) is a particularly intriguing system to explore the genetic basis of SI. Previous crossing experiments reported two distinct self-pollen recognition mechanisms operating across relatively short phylogenetic distance within the family: sporophytic SI in *Linanthus parviflorus* and gametophytic SI in *P. drummondii* (Levin 1993; Goodwillie 1997). Our crossing data contradicts previous reports of gametophytic SI in *P. drummondii* and conclusively demonstrates sporophytic control of self-pollen recognition. Prior crossing data was likely confounded by the presence of segregating SC variants, muddying patterns of cross-compatibility between related individuals. By simultaneously mapping the genetic basis of variation in the SI response, we were able to control for such segregating SC variants. Our findings are consistent with a single SI system operating in Polemoniaceae, present in both *Linanthus* and *Phlox*. Interestingly, gametophytic SI is known for multiple families related to Polemoniaceae, suggesting that an independent evolution of sporophytic SI occurred within the lineage containing *Phlox* (Igic *et al*. 2008). By mapping the genomic position of the *Phlox S*-locus and identifying regions containing genes potentially in the self-pollen rejection pathway, our work is a foundational step towards identifying the genetic basis of a novel homomorphic SI mechanism (Zhang *et al*. 2024; Ramanauskas *et al*. 2025).

## Data accessibility

Raw sequencing reads for all genotyped individuals and RNA sequencing are deposited in the NCBI short read archive (SUB15344310). The linkage map, phenotype-genotype table and BAM files used genetic mapping are available from the Dryad Digital Repository (doi:10.5061/dryad.7m0cfxq73).

## Authors’ contributions

G.A.B. and R.H. conceived of the study. All authors participated in data collection including field-collections, phenotyping, and sequencing. G.A.B. performed all data analyses and wrote the manuscript together with R.H.

## Acknowledgements

We thank Samridhi Chaturvedi for participating in seed-collection efforts, Felix Wu for their insights into linkage map construction, and Anna Feller for their guidance in QTL analyses. The Arnold Arboretum Greenhouse staff provided essential plant maintenance, and we would like to specifically acknowledge Lee Toomey, Megan Ardolino, Scott Pedemonte, and Mike Barrett for their *Phlox* care. Many thanks to the Bauer Sequencing Core for sequencing services and to the Harvard FAS Informatics Group for bioinformatics support. Comments from Boris Igić, Christina Steinecke, and three anonymous reviewers significantly improved our manuscript, and we appreciate their contributions.

