## Supplementary Figure 1 for "The genetic architecture of quantitative variation in the self-incompatibility response within *Phlox drummondii* (Polemoniaceae)"

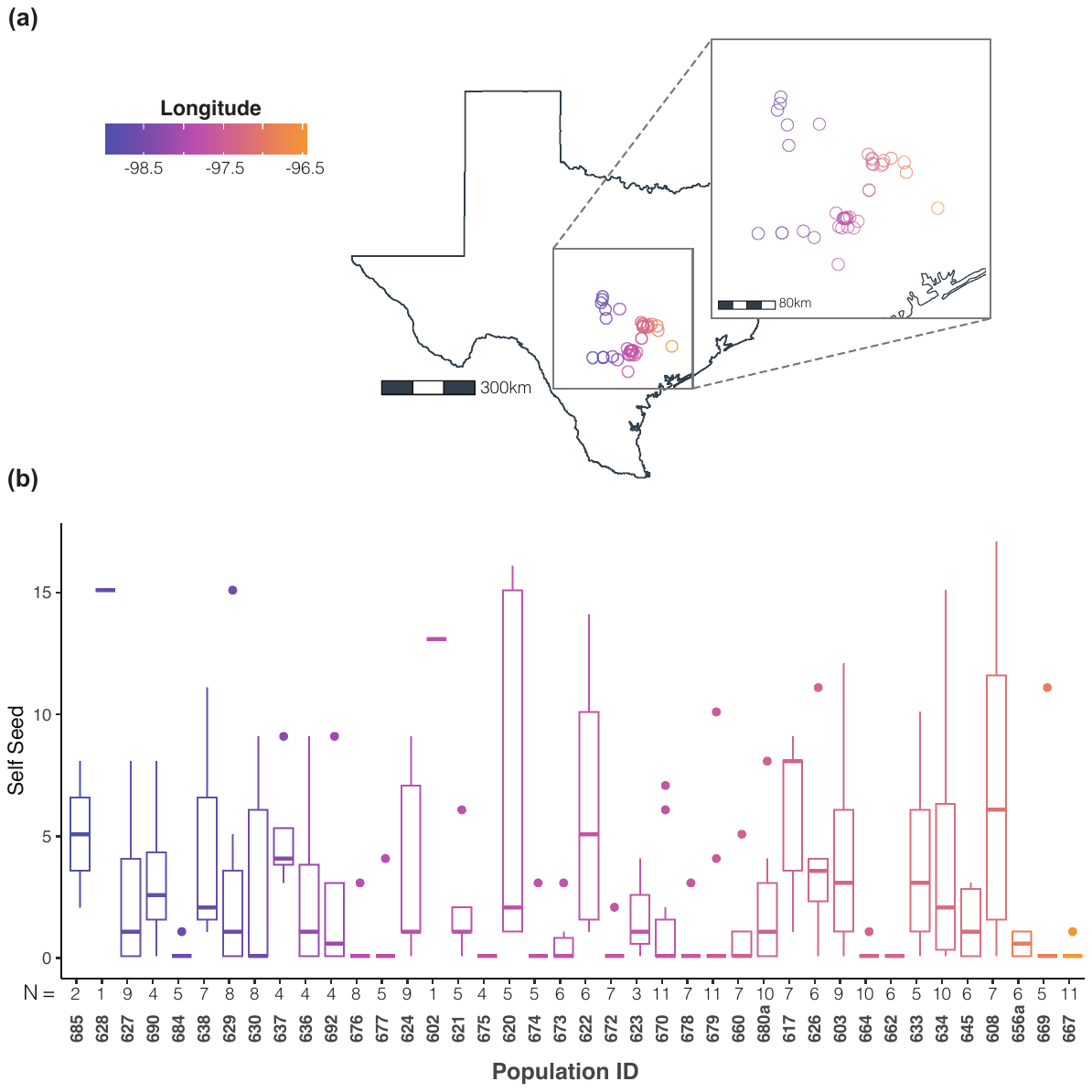

**Figure S1. (a)** Map indicating the locations of populations sampled. Points are colored by longitude. **(b)** The distribution of seed set following self-pollination for each population organized and colored by longitude. The number of individuals per population is listed directly below the x-axis.
