## Supplementary Figure 2 for "The genetic architecture of quantitative variation in the self-incompatibility response within *Phlox drummondii* (Polemoniaceae)"

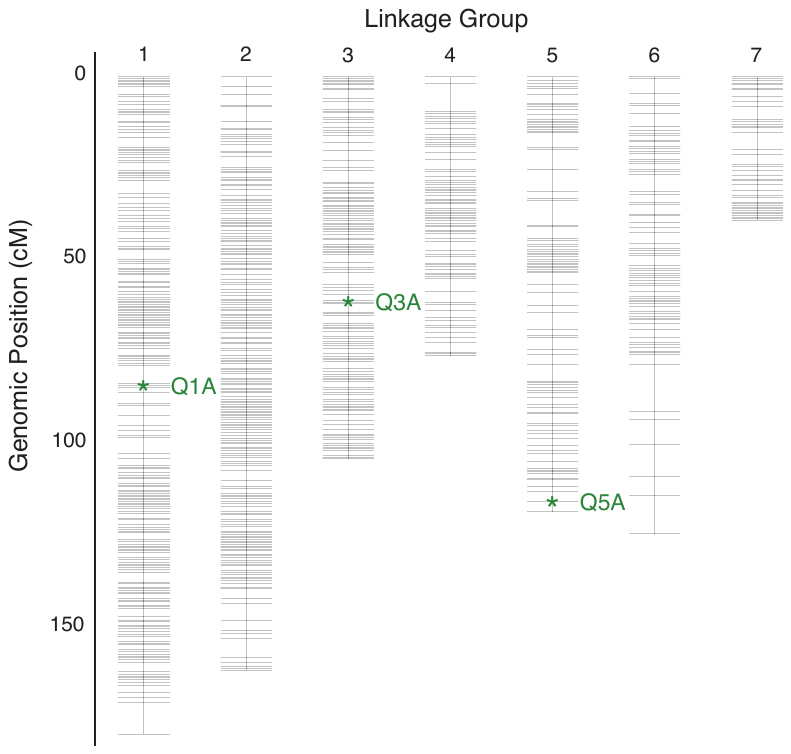

**Figure S2.** Family A linkage map generated from F2s using 6992 markers across seven linkage groups. Green stars indicate markers with the highest LOD score under significant QTL peaks in the Family A SI vs. SC cross.
