## Supplementary Figure 3 for "The genetic architecture of quantitative variation in the self-incompatibility response within *Phlox drummondii* (Polemoniaceae)"

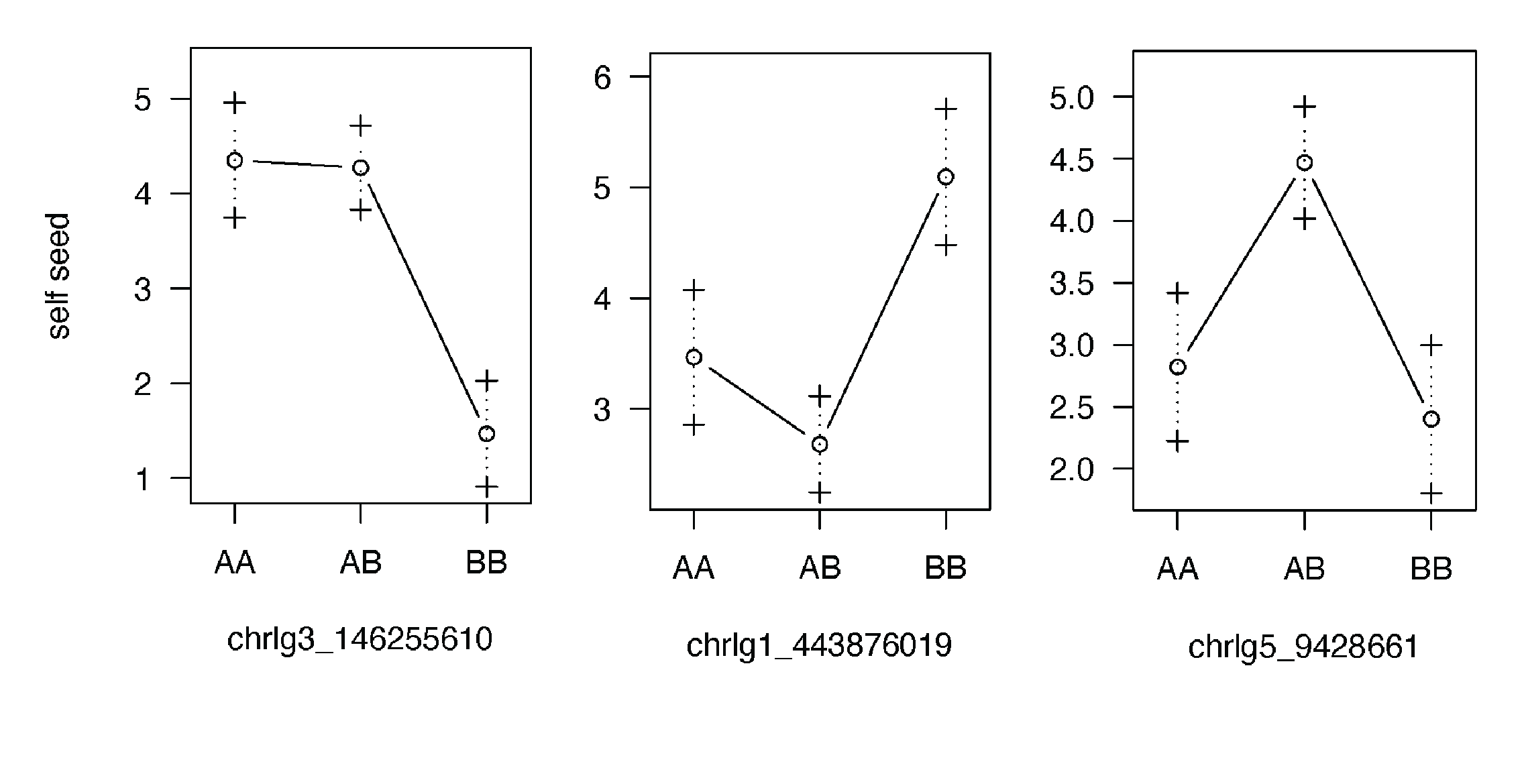

**Figure S3.** Effect plots at the markers with the highest LOD score under significant QTL peaks in the Family A SI vs. SC cross. The genotype “AA” indicates homozygous for the self-compatible parent allele (608-G1-2) while the genotype “BB” indicates homozygous for the self-incompatible parent allele (633-3-2).
