## Supplementary Figure 4 for "The genetic architecture of quantitative variation in the self-incompatibility response within *Phlox drummondii* (Polemoniaceae)"

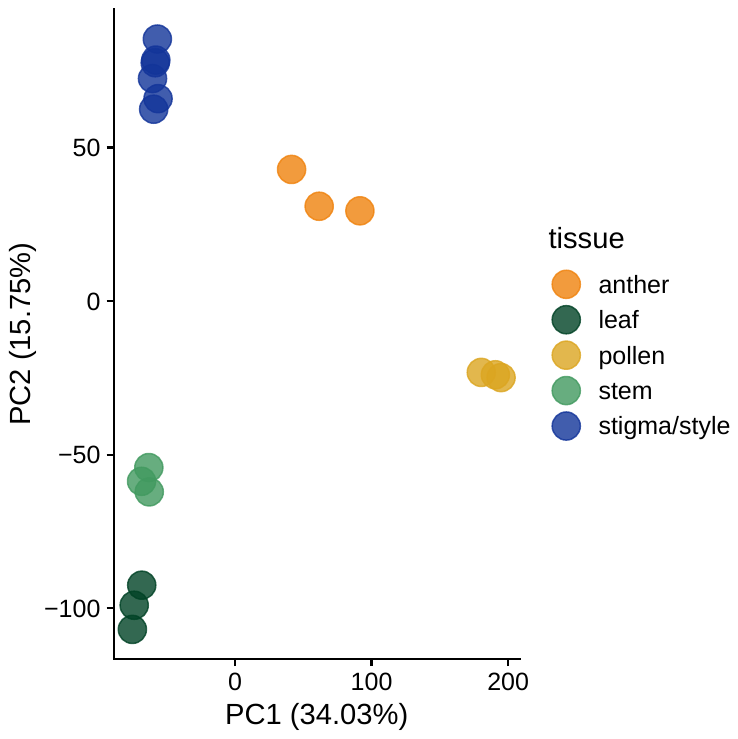

**Figure S4.** Principal component analysis (PCA) of gene expression across the five tissues sampled. Each point represents a unique sample and colors indicate tissue type.
