## Supplementary Figure 5 for "The genetic architecture of quantitative variation in the self-incompatibility response within *Phlox drummondii* (Polemoniaceae)"

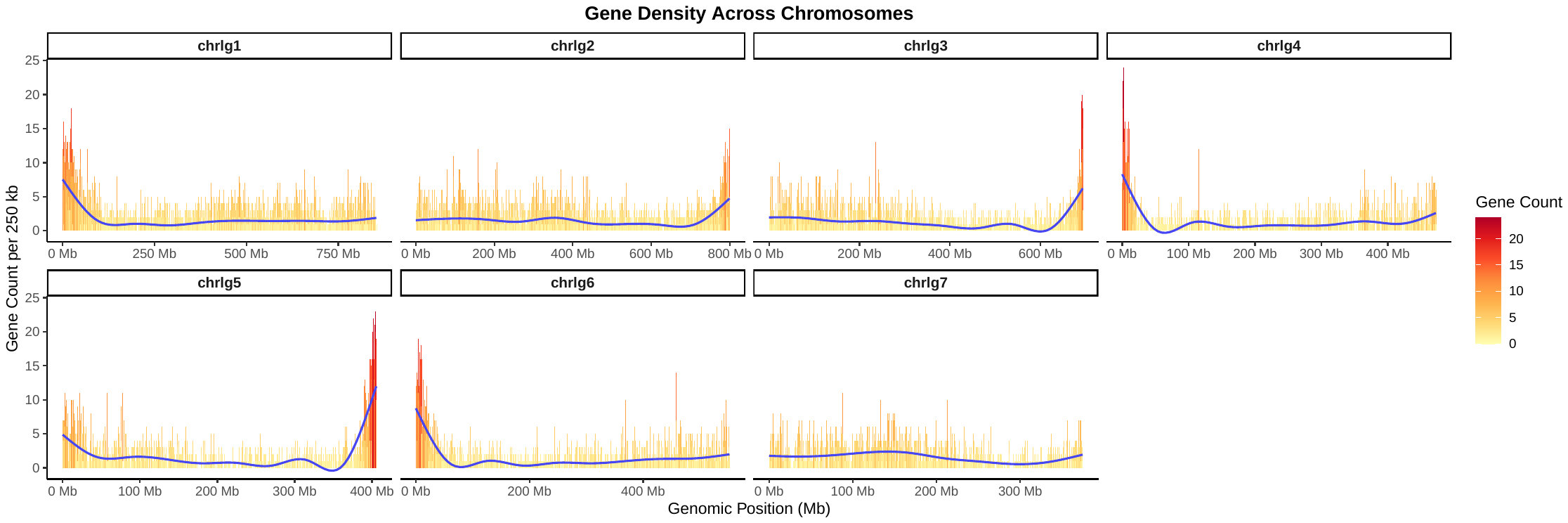

**Figure S5.** Gene density per 250kb window plotted against genomic position in *Phlox drummondii*. Bars are color scaled by gene count to emphasize relative density across windows. A smoothed line (GAM, cubic spline basis) is plotted in blue.
