## Supplementary Table 1 for "The genetic architecture of quantitative variation in the self-incompatibility response within *Phlox drummondii* (Polemoniaceae)"

| **Table S1.** Field-collected individuals used to assess quantitative variation in the self-incompatibility response | | | | | | | | |  |
| --- | --- | --- | --- | --- | --- | --- | --- | --- | --- |
|  |  |  |  |  | **Number of Seeds** | | **Number of Flowers** | |  |
| **Individual** | **Population** | **Latitude** | **Longitude** | **Year Grown** | **Self** | **Outcross** | **Self** | **Outcross** |  |
| 29 | 677 | 28.842167 | -97.845583 | 2020 | 0 | 16 | 8 | 8 |  |
| 57 | 677 | 28.842167 | -97.845583 | 2020 | 0 | 17 | 8 | 8 |  |
| 77 | 677 | 28.842167 | -97.845583 | 2020 | 0 | 17 | 8 | 8 |  |
| 123 | 677 | 28.842167 | -97.845583 | 2020 | 0 | 21 | 8 | 8 |  |
| 146 | 677 | 28.842167 | -97.845583 | 2020 | 4 | 23 | 8 | 8 |  |
| 1192 | 636 | 29.16865 | -98.18208 | 2014 | 0 | 19 | 8 | 6 |  |
| 636-33-1 | 636 | 29.16868 | -98.18208 | 2015 | 2 | 22 | 8 | 8 |  |
| 636-9-1 | 636 | 29.16868 | -98.18208 | 2015 | 9 | 23 | 8 | 8 |  |
| 636-9-2 | 636 | 29.16868 | -98.18208 | 2015 | 0 | 24 | 8 | 8 |  |
| 126 | 685 | 29.218909 | -98.972419 | 2020 | 2 | 20 | 8 | 8 |  |
| 257 | 685 | 29.218909 | -98.972419 | 2020 | 8 | 23 | 8 | 8 |  |
| 2 | 684 | 29.22645 | -98.637083 | 2020 | 0 | 23 | 8 | 8 |  |
| 115 | 684 | 29.22645 | -98.637083 | 2020 | 1 | 23 | 8 | 8 |  |
| 127 | 684 | 29.22645 | -98.637083 | 2020 | 0 | 19 | 8 | 8 |  |
| 183 | 684 | 29.22645 | -98.637083 | 2020 | 0 | 24 | 8 | 8 |  |
| 224 | 684 | 29.22645 | -98.637083 | 2020 | 0 | 20 | 8 | 8 |  |
| 638-24-2 | 638 | 29.22645 | -98.63708 | 2015 | 1 | 17 | 8 | 8 |  |
| 638-21-1 | 638 | 29.22645 | -98.63708 | 2015 | 1 | 18 | 8 | 8 |  |
| 638-5-1 | 638 | 29.22645 | -98.63708 | 2015 | 11 | 18 | 8 | 8 |  |
| 638-15-3 | 638 | 29.22645 | -98.63708 | 2015 | 7 | 19 | 8 | 8 |  |
| 638-7-1 | 638 | 29.22645 | -98.63708 | 2015 | 2 | 19 | 8 | 8 |  |
| 638-31-1 | 638 | 29.22645 | -98.63708 | 2015 | 2 | 20 | 8 | 8 |  |
| 638-24-1 | 638 | 29.22645 | -98.63708 | 2015 | 6 | 25 | 8 | 8 |  |
| 637-15-1 | 637 | 29.24847 | -98.33517 | 2015 | 4 | 18 | 8 | 8 |  |
| 637-18-1 | 637 | 29.24847 | -98.33517 | 2015 | 9 | 18 | 8 | 8 |  |
| 637-12-2 | 637 | 29.24847 | -98.33517 | 2015 | 3 | 20 | 8 | 8 |  |
| 1145 | 637 | 29.24847 | -98.33517 | 2014 | 4 | 21 | 8 | 9 |  |
| 7 | 678 | 29.2864167 | -97.6241 | 2020 | 0 | 22 | 8 | 8 |  |
| 38 | 678 | 29.2864167 | -97.6241 | 2020 | 3 | 23 | 8 | 8 |  |
| 81 | 678 | 29.2864167 | -97.6241 | 2020 | 0 | 23 | 8 | 8 |  |
| 109 | 678 | 29.2864167 | -97.6241 | 2020 | 0 | 24 | 8 | 8 |  |
| 143 | 678 | 29.2864167 | -97.6241 | 2020 | 0 | 21 | 8 | 8 |  |
| 160 | 678 | 29.2864167 | -97.6241 | 2020 | 0 | 24 | 8 | 8 |  |
| 215 | 678 | 29.2864167 | -97.6241 | 2020 | 0 | 22 | 8 | 8 |  |
| 1146 | 602 | 29.28728 | -97.79493 | 2014 | 13 | 18 | 8 | 6 |  |
| 623-9-1 | 623 | 29.29643 | -97.7084 | 2015 | 0 | 17 | 8 | 8 |  |
| 623-14-1 | 623 | 29.29643 | -97.7084 | 2015 | 4 | 20 | 8 | 8 |  |
| 623-17-1 | 623 | 29.29643 | -97.7084 | 2015 | 1 | 24 | 8 | 8 |  |
| 624-8-1 | 624 | 29.30333 | -97.83595 | 2015 | 5 | 19 | 8 | 8 |  |
| 624-13-1 | 624 | 29.30333 | -97.83595 | 2015 | 1 | 21 | 8 | 8 |  |
| 624-12-2 | 624 | 29.30333 | -97.83595 | 2015 | 7 | 22 | 8 | 8 |  |
| 624-32-1 | 624 | 29.30333 | -97.83595 | 2015 | 7 | 25 | 8 | 8 |  |
| 51 | 624 | 29.303333 | -97.83595 | 2020 | 1 | 20 | 8 | 8 |  |
| 68 | 624 | 29.303333 | -97.83595 | 2020 | 0 | 21 | 8 | 8 |  |
| 78 | 624 | 29.303333 | -97.83595 | 2020 | 9 | 21 | 8 | 8 |  |
| 132 | 624 | 29.303333 | -97.83595 | 2020 | 1 | 19 | 8 | 8 |  |
| 188 | 624 | 29.303333 | -97.83595 | 2020 | 0 | 18 | 8 | 8 |  |
| 37 | 679 | 29.367483 | -97.566617 | 2020 | 0 | 21 | 8 | 8 |  |
| 60 | 679 | 29.367483 | -97.566617 | 2020 | 0 | 19 | 8 | 8 |  |
| 82 | 679 | 29.367483 | -97.566617 | 2020 | 0 | 23 | 8 | 8 |  |
| 97 | 679 | 29.367483 | -97.566617 | 2020 | 0 | 25 | 8 | 8 |  |
| 119 | 679 | 29.367483 | -97.566617 | 2020 | 0 | 16 | 8 | 8 |  |
| 121 | 679 | 29.367483 | -97.566617 | 2020 | 4 | 23 | 8 | 8 |  |
| 158 | 679 | 29.367483 | -97.566617 | 2020 | 10 | 21 | 8 | 8 |  |
| 174 | 679 | 29.367483 | -97.566617 | 2020 | 0 | 24 | 8 | 8 |  |
| 197 | 679 | 29.367483 | -97.566617 | 2020 | 0 | 20 | 8 | 8 |  |
| 226 | 679 | 29.367483 | -97.566617 | 2020 | 0 | 24 | 8 | 8 |  |
| 229 | 679 | 29.367483 | -97.566617 | 2020 | 0 | 21 | 8 | 8 |  |
| 622-25-1 | 622 | 29.39867 | -97.72707 | 2015 | 3 | 18 | 8 | 8 |  |
| 622-9-1 | 622 | 29.39867 | -97.72707 | 2015 | 7 | 18 | 8 | 8 |  |
| 622-22-1 | 622 | 29.39867 | -97.72707 | 2015 | 1 | 20 | 8 | 8 |  |
| 622-3-2 | 622 | 29.39867 | -97.72707 | 2015 | 1 | 21 | 8 | 8 |  |
| 622-3-3 | 622 | 29.39867 | -97.72707 | 2015 | 14 | 21 | 8 | 8 |  |
| 622-8-2 | 622 | 29.39867 | -97.72707 | 2015 | 11 | 21 | 8 | 8 |  |
| 101 | 673 | 29.3992167 | -97.746583 | 2020 | 3 | 22 | 8 | 8 |  |
| 105 | 673 | 29.3992167 | -97.746583 | 2020 | 1 | 20 | 8 | 8 |  |
| 180 | 673 | 29.3992167 | -97.746583 | 2020 | 0 | 23 | 8 | 8 |  |
| 186 | 673 | 29.3992167 | -97.746583 | 2020 | 0 | 18 | 8 | 8 |  |
| 217 | 673 | 29.3992167 | -97.746583 | 2020 | 0 | 18 | 8 | 8 |  |
| 220 | 673 | 29.3992167 | -97.746583 | 2020 | 0 | 19 | 8 | 8 |  |
| 621-13-2 | 621 | 29.4024 | -97.77452 | 2015 | 0 | 16 | 8 | 8 |  |
| 621-12-1 | 621 | 29.4024 | -97.77452 | 2015 | 6 | 18 | 8 | 8 |  |
| 621-24-3 | 621 | 29.4024 | -97.77452 | 2015 | 2 | 19 | 8 | 8 |  |
| 621-14-1 | 621 | 29.4024 | -97.77452 | 2015 | 1 | 20 | 8 | 8 |  |
| 621-24-1 | 621 | 29.4024 | -97.77452 | 2015 | 1 | 22 | 8 | 8 |  |
| 35 | 675 | 29.4024 | -97.774517 | 2020 | 0 | 22 | 8 | 8 |  |
| 113 | 675 | 29.4024 | -97.774517 | 2020 | 0 | 17 | 8 | 8 |  |
| 206 | 675 | 29.4024 | -97.774517 | 2020 | 0 | 21 | 8 | 8 |  |
| 235 | 675 | 29.4024 | -97.774517 | 2020 | 0 | 20 | 8 | 8 |  |
| 620-4-2 | 620 | 29.40755 | -97.75622 | 2015 | 1 | 18 | 8 | 8 |  |
| 620-6-1 | 620 | 29.40755 | -97.75622 | 2015 | 1 | 19 | 8 | 8 |  |
| 620-17-1 | 620 | 29.40755 | -97.75622 | 2015 | 16 | 20 | 8 | 8 |  |
| 620-13-1 | 620 | 29.40755 | -97.75622 | 2015 | 2 | 23 | 8 | 8 |  |
| 620-10-1 | 620 | 29.40755 | -97.75622 | 2015 | 15 | 24 | 8 | 8 |  |
| 22 | 674 | 29.40755 | -97.756217 | 2020 | 0 | 16 | 8 | 8 |  |
| 40 | 674 | 29.40755 | -97.756217 | 2020 | 0 | 17 | 8 | 8 |  |
| 65 | 674 | 29.40755 | -97.756217 | 2020 | 3 | 20 | 8 | 8 |  |
| 94 | 674 | 29.40755 | -97.756217 | 2020 | 0 | 21 | 8 | 8 |  |
| 200 | 674 | 29.40755 | -97.756217 | 2020 | 0 | 18 | 8 | 8 |  |
| 31 | 672 | 29.4163333 | -97.719117 | 2020 | 0 | 19 | 8 | 8 |  |
| 50 | 672 | 29.4163333 | -97.719117 | 2020 | 0 | 21 | 8 | 8 |  |
| 111 | 672 | 29.4163333 | -97.719117 | 2020 | 2 | 23 | 8 | 8 |  |
| 120 | 672 | 29.4163333 | -97.719117 | 2020 | 0 | 17 | 8 | 8 |  |
| 134 | 672 | 29.4163333 | -97.719117 | 2020 | 0 | 21 | 8 | 8 |  |
| 159 | 672 | 29.4163333 | -97.719117 | 2020 | 0 | 23 | 8 | 8 |  |
| 211 | 672 | 29.4163333 | -97.719117 | 2020 | 0 | 18 | 8 | 8 |  |
| 9 | 670 | 29.4174167 | -97.6808 | 2020 | 6 | 24 | 8 | 8 |  |
| 10 | 670 | 29.4174167 | -97.6808 | 2020 | 0 | 24 | 8 | 8 |  |
| 23 | 670 | 29.4174167 | -97.6808 | 2020 | 0 | 24 | 8 | 8 |  |
| 63 | 670 | 29.4174167 | -97.6808 | 2020 | 2 | 20 | 8 | 8 |  |
| 70 | 670 | 29.4174167 | -97.6808 | 2020 | 1 | 20 | 8 | 8 |  |
| 124 | 670 | 29.4174167 | -97.6808 | 2020 | 0 | 20 | 8 | 8 |  |
| 154 | 670 | 29.4174167 | -97.6808 | 2020 | 0 | 21 | 8 | 8 |  |
| 171 | 670 | 29.4174167 | -97.6808 | 2020 | 7 | 18 | 8 | 8 |  |
| 185 | 670 | 29.4174167 | -97.6808 | 2020 | 0 | 24 | 8 | 8 |  |
| 203 | 670 | 29.4174167 | -97.6808 | 2020 | 0 | 16 | 8 | 8 |  |
| 233 | 670 | 29.4174167 | -97.6808 | 2020 | 0 | 18 | 8 | 8 |  |
| 1 | 676 | 29.4689024 | -97.873746 | 2020 | 0 | 22 | 8 | 8 |  |
| 19 | 676 | 29.4689024 | -97.873746 | 2020 | 0 | 22 | 8 | 8 |  |
| 49 | 676 | 29.4689024 | -97.873746 | 2020 | 3 | 25 | 8 | 8 |  |
| 83 | 676 | 29.4689024 | -97.873746 | 2020 | 0 | 18 | 8 | 8 |  |
| 88 | 676 | 29.4689024 | -97.873746 | 2020 | 0 | 24 | 8 | 8 |  |
| 173 | 676 | 29.4689024 | -97.873746 | 2020 | 0 | 18 | 8 | 8 |  |
| 179 | 676 | 29.4689024 | -97.873746 | 2020 | 0 | 18 | 8 | 8 |  |
| 194 | 676 | 29.4689024 | -97.873746 | 2020 | 0 | 20 | 8 | 8 |  |
| 4 | 667 | 29.528817 | -96.441233 | 2020 | 0 | 23 | 8 | 8 |  |
| 6 | 667 | 29.528817 | -96.441233 | 2020 | 0 | 24 | 8 | 8 |  |
| 13 | 667 | 29.528817 | -96.441233 | 2020 | 0 | 24 | 8 | 8 |  |
| 15 | 667 | 29.528817 | -96.441233 | 2020 | 0 | 22 | 8 | 8 |  |
| 201 | 667 | 29.528817 | -96.441233 | 2020 | 0 | 19 | 8 | 8 |  |
| 254 | 667 | 29.528817 | -96.441233 | 2020 | 0 | 21 | 8 | 8 |  |
| 260 | 667 | 29.528817 | -96.441233 | 2020 | 0 | 16 | 8 | 8 |  |
| 264 | 667 | 29.528817 | -96.441233 | 2020 | 0 | 21 | 8 | 8 | Family C |
| 273 | 667 | 29.528817 | -96.441233 | 2020 | 0 | 20 | 8 | 8 |  |
| 278 | 667 | 29.528817 | -96.441233 | 2020 | 1 | 22 | 8 | 8 |  |
| 281 | 667 | 29.528817 | -96.441233 | 2020 | 0 | 20 | 8 | 8 |  |
| 62 | 680a | 29.7466167 | -97.4097 | 2020 | 3 | 21 | 8 | 8 |  |
| 76 | 680a | 29.7466167 | -97.4097 | 2020 | 4 | 23 | 8 | 8 |  |
| 93 | 680a | 29.7466167 | -97.4097 | 2020 | 0 | 23 | 8 | 8 |  |
| 130 | 680a | 29.7466167 | -97.4097 | 2020 | 0 | 18 | 8 | 8 |  |
| 133 | 680a | 29.7466167 | -97.4097 | 2020 | 8 | 16 | 8 | 8 |  |
| 198 | 680a | 29.7466167 | -97.4097 | 2020 | 0 | 24 | 8 | 8 |  |
| 205 | 680a | 29.7466167 | -97.4097 | 2020 | 1 | 23 | 8 | 8 |  |
| 219 | 680a | 29.7466167 | -97.4097 | 2020 | 3 | 17 | 8 | 8 |  |
| 222 | 680a | 29.7466167 | -97.4097 | 2020 | 0 | 23 | 8 | 8 |  |
| 230 | 680a | 29.7466167 | -97.4097 | 2020 | 1 | 19 | 8 | 8 |  |
| 617-9-1 | 617 | 29.75098 | -97.40962 | 2015 | 9 | 16 | 8 | 8 |  |
| 617-32-1 | 617 | 29.75098 | -97.40962 | 2015 | 1 | 17 | 8 | 8 |  |
| 617-10-1 | 617 | 29.75098 | -97.40962 | 2015 | 5 | 21 | 8 | 8 |  |
| 617-5-1 | 617 | 29.75098 | -97.40962 | 2015 | 8 | 22 | 8 | 8 |  |
| 617-8-4 | 617 | 29.75098 | -97.40962 | 2015 | 2 | 22 | 8 | 8 |  |
| 617-12-1 | 617 | 29.75098 | -97.40962 | 2015 | 8 | 23 | 8 | 8 |  |
| 617-15-5 | 617 | 29.75098 | -97.40962 | 2015 | 8 | 23 | 8 | 8 |  |
| 103 | 669 | 29.965443 | -96.885429 | 2020 | 11 | 18 | 8 | 8 |  |
| 129 | 669 | 29.965443 | -96.885429 | 2020 | 0 | 22 | 8 | 8 |  |
| 150 | 669 | 29.965443 | -96.885429 | 2020 | 0 | 20 | 8 | 8 |  |
| 162 | 669 | 29.965443 | -96.885429 | 2020 | 0 | 24 | 8 | 8 |  |
| 165 | 669 | 29.965443 | -96.885429 | 2020 | 0 | 23 | 8 | 8 |  |
| 633-18-1 | 633 | 30.0505 | -97.2378 | 2015 | 10 | 16 | 8 | 8 |  |
| 633-9-1 | 633 | 30.0505 | -97.2378 | 2015 | 3 | 16 | 8 | 8 |  |
| 633-18-2 | 633 | 30.0505 | -97.2378 | 2015 | 1 | 20 | 8 | 8 |  |
| 633-3-2 | 633 | 30.0505 | -97.2378 | 2015 | 0 | 24 | 8 | 8 | Family A |
| 633-4-1 | 633 | 30.0505 | -97.2378 | 2015 | 6 | 24 | 8 | 8 |  |
| 46 | 662 | 30.0621514 | -97.350295 | 2020 | 0 | 20 | 8 | 8 |  |
| 116 | 662 | 30.0621514 | -97.350295 | 2020 | 0 | 16 | 8 | 8 |  |
| 56 | 662 | 30.0665023 | -97.352609 | 2020 | 0 | 22 | 8 | 8 |  |
| 152 | 662 | 30.0665023 | -97.352609 | 2020 | 0 | 23 | 8 | 8 |  |
| 156 | 662 | 30.0665023 | -97.352609 | 2020 | 0 | 16 | 8 | 8 |  |
| 240 | 662 | 30.0665023 | -97.352609 | 2020 | 0 | 23 | 8 | 8 |  |
| 634-23-1 | 634 | 30.06688 | -97.21388 | 2015 | 1 | 16 | 8 | 8 |  |
| 634-28-1 | 634 | 30.06688 | -97.21388 | 2015 | 1 | 19 | 8 | 8 |  |
| 634-32-1 | 634 | 30.06688 | -97.21388 | 2015 | 10 | 22 | 8 | 8 |  |
| 634-17-1 | 634 | 30.06688 | -97.21388 | 2015 | 7 | 23 | 8 | 8 |  |
| 634-44-1 | 634 | 30.06688 | -97.21388 | 2015 | 4 | 23 | 8 | 8 |  |
| 11 | 634 | 30.066883 | -97.213883 | 2020 | 15 | 19 | 8 | 8 |  |
| 32 | 634 | 30.066883 | -97.213883 | 2020 | 0 | 22 | 8 | 8 |  |
| 102 | 634 | 30.066883 | -97.213883 | 2020 | 3 | 20 | 8 | 8 |  |
| 163 | 634 | 30.066883 | -97.213883 | 2020 | 0 | 24 | 8 | 8 |  |
| 170 | 634 | 30.066883 | -97.213883 | 2020 | 0 | 18 | 8 | 8 |  |
| 603-25-1 | 603 | 30.0704 | -97.36613 | 2015 | 7 | 16 | 8 | 8 |  |
| 603-21-1 | 603 | 30.0704 | -97.36613 | 2015 | 3 | 17 | 8 | 8 |  |
| 603-25-3 | 603 | 30.0704 | -97.36613 | 2015 | 6 | 18 | 8 | 8 |  |
| 603-6-1 | 603 | 30.0704 | -97.36613 | 2015 | 1 | 18 | 8 | 8 |  |
| 603-23-1 | 603 | 30.0704 | -97.36613 | 2015 | 5 | 19 | 8 | 8 |  |
| 603-26-1 | 603 | 30.0704 | -97.36613 | 2015 | 0 | 19 | 8 | 8 |  |
| 603-27-2 | 603 | 30.0704 | -97.36613 | 2015 | 1 | 19 | 8 | 8 |  |
| 603-27-1 | 603 | 30.0704 | -97.36613 | 2015 | 3 | 21 | 8 | 8 |  |
| 603-3-2 | 603 | 30.0704 | -97.36613 | 2015 | 12 | 21 | 8 | 8 |  |
| 1105 | 656a | 30.0852333 | -96.916167 | 2014 | 1 | 17 | 8 | 9 |  |
| 43 | 656a | 30.0852333 | -96.916167 | 2020 | 1 | 23 | 8 | 8 |  |
| 64 | 656a | 30.0852333 | -96.916167 | 2020 | 0 | 17 | 8 | 8 |  |
| 221 | 656a | 30.0852333 | -96.916167 | 2020 | 1 | 17 | 8 | 8 |  |
| 234 | 656a | 30.0852333 | -96.916167 | 2020 | 0 | 23 | 8 | 8 |  |
| 238 | 656a | 30.0852333 | -96.916167 | 2020 | 0 | 18 | 8 | 8 |  |
| 645-25-1 | 645 | 30.1049 | -97.20428 | 2015 | 3 | 16 | 8 | 8 |  |
| 645-18-1 | 645 | 30.1049 | -97.20428 | 2015 | 0 | 18 | 8 | 8 |  |
| 645-30-1 | 645 | 30.1049 | -97.20428 | 2015 | 0 | 18 | 8 | 8 |  |
| 645-4-2 | 645 | 30.1049 | -97.20428 | 2015 | 0 | 21 | 8 | 8 |  |
| 645-3-1 | 645 | 30.1049 | -97.20428 | 2015 | 3 | 22 | 8 | 8 |  |
| 645-12-1 | 645 | 30.1049 | -97.20428 | 2015 | 2 | 23 | 8 | 8 |  |
| 21 | 664 | 30.116 | -97.365 | 2020 | 0 | 23 | 8 | 8 |  |
| 131 | 664 | 30.116 | -97.365 | 2020 | 0 | 22 | 8 | 8 |  |
| 223 | 664 | 30.116 | -97.365 | 2020 | 0 | 18 | 8 | 8 |  |
| 242 | 664 | 30.116 | -97.365 | 2020 | 0 | 23 | 8 | 8 |  |
| 252 | 664 | 30.116 | -97.365 | 2020 | 0 | 21 | 8 | 8 |  |
| 253 | 664 | 30.116 | -97.365 | 2020 | 0 | 22 | 8 | 8 |  |
| 259 | 664 | 30.116 | -97.365 | 2020 | 0 | 23 | 8 | 8 | Family B |
| 267 | 664 | 30.116 | -97.365 | 2020 | 0 | 22 | 8 | 8 |  |
| 269 | 664 | 30.116 | -97.365 | 2020 | 0 | 20 | 8 | 8 |  |
| 271 | 664 | 30.116 | -97.365 | 2020 | 1 | 23 | 8 | 8 |  |
| 608-E1-1 | 608 | 30.13343 | -97.10107 | 2015 | 0 | 16 | 8 | 8 |  |
| 608-G1-3 | 608 | 30.13343 | -97.10107 | 2015 | 6 | 17 | 8 | 8 |  |
| 608-F1-1A | 608 | 30.13343 | -97.10107 | 2015 | 13 | 19 | 8 | 8 |  |
| 608-B1-1 | 608 | 30.13343 | -97.10107 | 2015 | 10 | 22 | 8 | 8 |  |
| 608-D1-1 | 608 | 30.13343 | -97.10107 | 2015 | 1 | 22 | 8 | 8 |  |
| 608-F3-1 | 608 | 30.13343 | -97.10107 | 2015 | 2 | 22 | 8 | 8 |  |
| 608-G1-2 | 608 | 30.13343 | -97.10107 | 2015 | 17 | 25 | 8 | 8 | Family A |
| 626-4-1 | 626 | 30.135 | -97.37135 | 2015 | 11 | 17 | 8 | 8 |  |
| 626-38-1 | 626 | 30.135 | -97.37135 | 2015 | 0 | 18 | 8 | 8 |  |
| 626-38-2 | 626 | 30.135 | -97.37135 | 2015 | 3 | 18 | 8 | 8 |  |
| 626-3-1 | 626 | 30.135 | -97.37135 | 2015 | 2 | 21 | 8 | 8 |  |
| 626-5-1 | 626 | 30.135 | -97.37135 | 2015 | 4 | 25 | 8 | 8 |  |
| 626-37-1 | 626 | 30.135 | -97.37135 | 2015 | 4 | 31 | 8 | 8 |  |
| 5 | 660 | 30.1803833 | -97.420717 | 2020 | 0 | 20 | 8 | 8 |  |
| 61 | 660 | 30.1803833 | -97.420717 | 2020 | 5 | 23 | 8 | 8 |  |
| 80 | 660 | 30.1803833 | -97.420717 | 2020 | 0 | 22 | 8 | 8 |  |
| 128 | 660 | 30.1803833 | -97.420717 | 2020 | 0 | 22 | 8 | 8 |  |
| 182 | 660 | 30.1803833 | -97.420717 | 2020 | 0 | 19 | 8 | 8 |  |
| 196 | 660 | 30.1803833 | -97.420717 | 2020 | 1 | 21 | 8 | 8 |  |
| 236 | 660 | 30.1803833 | -97.420717 | 2020 | 1 | 23 | 8 | 8 |  |
| 161 | 630 | 30.2901 | -98.540383 | 2020 | 0 | 18 | 8 | 8 |  |
| 42 | 630 | 30.2901 | -98.540383 | 2020 | 0 | 24 | 8 | 8 |  |
| 95 | 630 | 30.2901 | -98.540383 | 2020 | 0 | 22 | 8 | 8 |  |
| 99 | 630 | 30.2901 | -98.540383 | 2020 | 0 | 20 | 8 | 8 |  |
| 630-12-2 | 630 | 30.2901 | -98.54038 | 2015 | 2 | 16 | 8 | 8 |  |
| 630-18-1 | 630 | 30.2901 | -98.54038 | 2015 | 6 | 18 | 8 | 8 |  |
| 630-42-1 | 630 | 30.2901 | -98.54038 | 2015 | 9 | 18 | 8 | 8 |  |
| 630-12-1 | 630 | 30.2901 | -98.54038 | 2015 | 0 | 22 | 8 | 8 |  |
| 630-38-2 | 630 | 30.2901 | -98.54038 | 2015 | 6 | 22 | 8 | 8 |  |
| 26 | 629 | 30.54095 | -98.56135 | 2020 | 0 | 23 | 8 | 8 |  |
| 72 | 629 | 30.54095 | -98.56135 | 2020 | 5 | 19 | 8 | 8 |  |
| 151 | 629 | 30.54095 | -98.56135 | 2020 | 0 | 24 | 8 | 8 |  |
| 172 | 629 | 30.54095 | -98.56135 | 2020 | 0 | 21 | 8 | 8 |  |
| 195 | 629 | 30.54095 | -98.56135 | 2020 | 3 | 24 | 8 | 8 |  |
| 204 | 629 | 30.54095 | -98.56135 | 2020 | 0 | 23 | 8 | 8 |  |
| 629-9-1 | 629 | 30.54095 | -98.56135 | 2015 | 15 | 22 | 8 | 8 |  |
| 629-34-2 | 629 | 30.54095 | -98.56135 | 2015 | 2 | 23 | 8 | 8 |  |
| 14 | 692 | 30.5497 | -98.110483 | 2020 | 9 | 21 | 8 | 8 |  |
| 117 | 692 | 30.5497 | -98.110483 | 2020 | 0 | 17 | 8 | 8 |  |
| 190 | 692 | 30.5497 | -98.110483 | 2020 | 1 | 17 | 8 | 8 |  |
| 244 | 692 | 30.5497 | -98.110483 | 2020 | 0 | 20 | 8 | 8 |  |
| 628-25-3 | 628 | 30.72118 | -98.70118 | 2015 | 15 | 17 | 8 | 8 |  |
| 627-44-1 | 627 | 30.8018 | -98.66287 | 2015 | 8 | 18 | 8 | 8 |  |
| 627-44-3 | 627 | 30.8018 | -98.66287 | 2015 | 6 | 21 | 8 | 8 |  |
| 627-1-2 | 627 | 30.8018 | -98.66287 | 2015 | 4 | 22 | 8 | 8 |  |
| 627-1-4 | 627 | 30.8018 | -98.66287 | 2015 | 0 | 24 | 8 | 8 |  |
| 8 | 627 | 30.8018 | -98.662867 | 2020 | 0 | 20 | 8 | 8 |  |
| 34 | 627 | 30.8018 | -98.662867 | 2020 | 0 | 18 | 8 | 8 |  |
| 90 | 627 | 30.8018 | -98.662867 | 2020 | 0 | 18 | 8 | 8 |  |
| 96 | 627 | 30.8018 | -98.662867 | 2020 | 1 | 24 | 8 | 8 |  |
| 108 | 627 | 30.8018 | -98.662867 | 2020 | 1 | 20 | 8 | 8 |  |
| 255 | 690 | 30.880867 | -98.653667 | 2020 | 8 | 23 | 8 | 8 |  |
| 262 | 690 | 30.880867 | -98.653667 | 2020 | 2 | 23 | 8 | 8 |  |
| 265 | 690 | 30.880867 | -98.653667 | 2020 | 3 | 20 | 8 | 8 |  |
| 268 | 690 | 30.880867 | -98.653667 | 2020 | 0 | 21 | 8 | 8 |  |
| Note. Individuals are sorted by latitude and longitude. All individuals grown in 2014 and 2015 were previously published in Roda & Hopkins (2019). Individuals used as parents for the three genetic mapping populations presented in this study (Family A, Family B, Family C) are indicated to the right of the table. | | | | | | | | |  |
