## Supplementary Table 2 for "The genetic architecture of quantitative variation in the self-incompatibility response within *Phlox drummondii* (Polemoniaceae)"

| **Table S2.** Linkage map summary statistics | | | | |
| --- | --- | --- | --- | --- |
|  |  |  | **Spacing Between Markers (cM)** | |
| **Linkage Group** | **Number of Markers** | **Length (cM)** | **Average** | **Maximum** |
| 1 | 2425 | 177.4 | 0.1 | 8.6 |
| 2 | 1778 | 160.2 | 0.1 | 5.0 |
| 3 | 1092 | 103.1 | 0.1 | 3.1 |
| 4 | 722 | 75.2 | 0.1 | 7.6 |
| 5 | 453 | 117.3 | 0.3 | 6.8 |
| 6 | 321 | 123.4 | 0.4 | 12.6 |
| 7 | 201 | 38.7 | 0.2 | 4.4 |
| Overall | 6992 | 795.4 | 0.1 | 12.6 |
