## Supplementary Table 3 for "The genetic architecture of quantitative variation in the self-incompatibility response within *Phlox drummondii* (Polemoniaceae)"

| **Table S3.** Summary of sequencing for bulk segregant analyses | | | | |
| --- | --- | --- | --- | --- |
|  |  |  | **Sequenced reads** | |
| **Comparison** | **Family** | **Bulk** | **Total** | **Aligned and mapped** |
| Variation in  Self-Incompatibility  (SI vs. SC) | B | SI (3,4,5) | 17,854,455 | 17,433,220 |
|  | B | SC (9) | 4,578,964 | 4,476,142 |
|  | C | SI (1,2,6) | 14,556,349 | 14,122,165 |
|  | C | SC (7) | 3,238,585 | 3,144,589 |
| S-locus Genotype  (S1S1 vs. S2S2 or  S3S3 vs. S4S4) | B | S1S1 (3) | 3,863,409 | 3,762,538 |
|  | B | S2S2 (4) | 6,142,016 | 6,004,851 |
|  | C | S3S3 (1) | 4,821,212 | 4,668,845 |
|  | C | S4S4 (2) | 9,726,467 | 9,444,952 |

Note. Bulks are listed by phenotype (self-incompatibility status or inferred S-locus genotype) with bulk number assigned for sequencing given in parentheses.
