## Supplementary Table 4 for "The genetic architecture of quantitative variation in the self-incompatibility response within *Phlox drummondii* (Polemoniaceae)"

| **QTL**  Table S4. Stigma/style expressed candidate genes in QTL intervals. Gene IDs sourced from *Phlox drummondii* annotation. | **Linkage Group** | **5' CDS position** | ***P. drummondii* Annotation ID** | **TAIR ID** | **e-value** | **Identity (%)** | **Length (bp)** | **Arabidopsis Homolog Description** |
| --- | --- | --- | --- | --- | --- | --- | --- | --- |
| Q3A | 3 | 600455494 | PD.60436 | AT1G25330 | 1e-61 | 82 | 239 | CESTA transcription factor, positive regulator of brassinosteroid biosynthesis |
| Q3A | 3 | 655042297 | PD.60922 | AT5G38450 | 5e-05 | 83 | 47 | CYP735A1, cytochrome P450 |
| Q3A | 3 | 202932972 | PD.56651 | AT2G19090 | 2e-08 | 71 | 136 | DUF630 family protein |
| Q3A | 3 | 458598066 | PD.59425 | AT5G59090 | 1e-05 | 68 | 282 | subtilase, serine-protease |
| Q3A | 3 | 263983747 | PD.57615 | AT5G55180 | 5e-128 | 68 | 1299 | O-Glycosyl hydrolases family 17 protein |
| Q3A | 3 | 634904848 | PD.60704 | AT5G42920 | 0.046 | 92 | 26 | encodes component of the putative THO/TREX complex, mRNA precursor transport |
| Q3A | 3 | 146782401 | PD.55820 | AT2G26530 | 4e-06 | 74 | 84 | Pheromone receptor-like protein involved in early elicitor signaling events, activation of MPKs and the formation of ROS |
| Q3A | 3 | 640260063 | PD.60756 | AT1G13550 | 3e-16 | 74 | 138 | hypothetical protein, DUF1626 |
| Q1A | 1 | 590691606 | PD.36744 | AT2G17480 | 7e-79 | 78 | 364 | MLO protein family, transmembrane protein |
| Q1A | 1 | 590704412 | PD.36747 | AT5G65970 | 1e-36 | 69 | 380 | MLO protein family, transmembrane protein |
| Q1A | 1 | 592838225 | PD.36790 | AT5G37020 | 0 | 76 | 1391 | Auxin response factor family |
| Q1A | 1 | 640868042 | PD.37530 | AT4G27290 | 8e-05 | 66 | 207 | S-locus lectin protein kinase family protein |
| Q1A | 1 | 613130547 | PD.37092 | AT5G65170 | 4e-12 | 76 | 103 | VQ motif-containing protein |
| Q1A | 1 | 578967031 | PD.36531 | AT3G14780 | 2e-04 | 85 | 46 | callose synthase |
| Q1A | 1 | 627447921 | PD.37301 | AT5G09750 | 2e-58 | 76 | 305 | bHLH transcription factor involved in transmitting tract/stigma development |
| Q1A | 1 | 588851215 | PD.36719 | no hit |  |  |  |  |
| Q1A | 1 | 566777233 | PD.36347 | AT5G66560 | 2e-51 | 74 | 331 | phototropic-responsive NPH3 family protein |
| Q1A | 1 | 654178686 | PD.37795 | AT5G03380 | 3e-04 | 68 | 118 | heavy metal transport/detoxification superfamily |
| Q1A | 1 | 648790039 | PD.37701 | AT2G22590 | 0.001 | 77 | 60 | UDP-Glycosyltransferase superfamily protein |
| S-locus | 2 | 770195889 | PD.51830 | AT1G74650 | 7e-67 | 76 | 361 | MYB31, wax regulator associated with reproductive development |
| S-locus & Q2C | 2 | 788934003 | PD.52568 | AT4G21380 | 5e-44 | 69 | 480 | S-locus lectin receptor-like kinase family |
| S-locus & Q2C | 2 | 735828363 | PD.51265 | AT1G77810 | 4e-94 | 78 | 436 | galactosyltransferase family protein |
| S-locus & Q2C | 2 | 791268733 | PD.52651 | AT5G37830 | 0.22 | 85 | 33 | 5-oxoprolinase that acts in the gluthione degradation pathway, oxoproline metabolism |
| S-locus & Q2C | 2 | 791717565 | PD.52670 | AT4G11280 | 3e-118 | 68 | 1232 | encodes a member of the 1-aminocyclopropane-1-carboxylate (ACC) synthase gene family |
| S-locus & Q2C | 2 | 791263774 | PD.52649 | AT1G14850 | 0.76 | 92 | 24 | encodes a protein similar to nucleoporin |
| S-locus & Q2C | 2 | 789511151 | PD.52576 | AT2G42350 | 9e-08 | 69 | 147 | RING/U-box superfamily protein |
| S-locus & Q2C | 2 | 785852525 | PD.52359 | AT5G51950 | 5e-30 | 76 | 181 | glucose-methanol-choline oxireductase family |
| Q5A | 5 | 115972118 | PD.70740 | AT1G75520 | 1e-37 | 86 | 132 | SHI gene family, positive regulator of photomorphogenesis |
| Q5A | 5 | 50246593 | PD.69752 | AT2G45190 | 1e-80 | 72 | 600 | YABBY family of trascriptional regulators involved in abaxial cell type specification |
| Q5A | 5 | 22170316 | PD.69276 | AT2G44260 | 2e-70 | 67 | 912 | DUF946 family protein |
| Q5A | 5 | 57029317 | PD.69863 | AT4G12110 | 4e-81 | 72 | 591 | SMO1 family of sterol 4alpha-methyl oxidases |
| Q5A | 5 | 123043753 | PD.70852 | AT4G37070 | 8e-12 | 80 | 79 | patatin-related phospholipase |
| Q5A & Q5B | 5 | 306727262 | PD.72277 | AT3G15540 | 9e-51 | 73 | 358 | primary auxin-responsive gene |
| Q5A & Q5B | 5 | 358528190 | PD.72635 | AT5G66460 | 3e-143 | 70 | 1168 | endo-beta-mannanase |
| Q5A & Q5B | 5 | 303347622 | PD.72244 | AT1G15540 | 2e-04 | 87 | 39 | 2-oxoglutarate-dependent dioxygenase-like |
| Q5B | 5 | 400138907 | PD.73874 | AT5G08590 | 0 | 77 | 1034 | member of SNF1-related protein kinases, activated by osmotic stress |
| Q5B | 5 | 400776598 | PD.73932 | AT1G23030 | 1e-07 | 72 | 105 | plant U-Box protein capable of acting as an E3 ligase, regulates environmental responses via selective degredation of targets |
| Q5B | 5 | 397674738 | PD.73638 | AT1G76600 | 6e-21 | 74 | 168 | PADRE protein |
| Q5B | 5 | 398405246 | PD.73720 | AT3G56490 | 1e-110 | 82 | 400 | adenylylsulfate sulfohyrdolase activity |
| Q4B | 4 | 240552498 | PD.65627 | AT1G53820 | 4e-18 | 73 | 154 | RING/U-box superfamily protein |
| Q4B  Note. TAIR ID is given for the *Arabidospis thaliana* homolog with the lowest E-Value following BLAST search in the TAIR database (arabidopsis.org) | 4 | 319312764 | PD.66449 | AT4G11410 | 9e-44 | 68 | 532 | NAD(P)-binding Rossmann-fold superfamily |
