## Supplementary Table 5 for "The genetic architecture of quantitative variation in the self-incompatibility response within *Phlox drummondii* (Polemoniaceae)"

Table S5. Normalized count data for stigma/style expressed candidate genes.

| ***P. drummondii* Annotation ID** | **Stigma and Style** | | | | | | **Anther** | | | **Leaf** | | | **Pollen** | | | **Stem** | | |
| --- | --- | --- | --- | --- | --- | --- | --- | --- | --- | --- | --- | --- | --- | --- | --- | --- | --- | --- |
|  | **1** | **2** | **3** | **4** | **5** | **6** | **1** | **2** | **3** | **1** | **2** | **3** | **1** | **2** | **3** | **1** | **2** | **3** |
| PD.60436 | 29481 | 36180 | 64202 | 56094 | 59299 | 65650 | 0 | 0 | 0 | 0 | 0 | 0 | 0 | 0 | 0 | 0 | 1 | 0 |
| PD.60922 | 53198 | 30631 | 79825 | 73152 | 93279 | 86148 | 579 | 161 | 397 | 510 | 433 | 318 | 0 | 0 | 0 | 241 | 150 | 159 |
| PD.56651 | 881 | 458 | 1888 | 3828 | 1968 | 2101 | 0 | 9 | 6 | 12 | 0 | 2 | 0 | 0 | 0 | 7 | 38 | 19 |
| PD.59425 | 8234 | 4222 | 30538 | 46210 | 20730 | 19815 | 0 | 0 | 0 | 1 | 2 | 14 | 0 | 0 | 0 | 8 | 6 | 0 |
| PD.57615 | 39845 | 32576 | 106074 | 136753 | 81506 | 101044 | 250 | 460 | 94 | 278 | 513 | 315 | 26 | 1 | 2 | 1738 | 2708 | 1702 |
| PD.60704 | 899 | 4483 | 2848 | 7884 | 2267 | 3846 | 0 | 0 | 0 | 11 | 12 | 9 | 0 | 0 | 0 | 5 | 42 | 70 |
| PD.55820 | 24 | 377 | 300 | 87 | 645 | 353 | 2 | 1 | 3 | 3 | 1 | 3 | 1 | 1 | 0 | 2 | 8 | 7 |
| PD.60756 | 1261 | 586 | 3077 | 163 | 2436 | 1366 | 10 | 6 | 4 | 7 | 2 | 0 | 16 | 3 | 1 | 10 | 7 | 0 |
| PD.36744 | 3719 | 2056 | 2981 | 4421 | 4726 | 4552 | 0 | 4 | 0 | 6 | 5 | 4 | 1 | 0 | 0 | 211 | 150 | 165 |
| PD.36747 | 25624 | 21256 | 63952 | 39101 | 35627 | 62822 | 29 | 50 | 9 | 427 | 376 | 239 | 0 | 0 | 5 | 388 | 316 | 260 |
| PD.36790 | 6822 | 4138 | 20709 | 22143 | 14756 | 18929 | 143 | 183 | 74 | 130 | 148 | 135 | 6 | 6 | 8 | 2149 | 2391 | 2201 |
| PD.37530 | 51151 | 71581 | 55314 | 120456 | 74447 | 103153 | 206 | 84 | 26 | 0 | 0 | 0 | 0 | 0 | 0 | 0 | 0 | 0 |
| PD.37092 | 1136 | 1237 | 3379 | 5437 | 1808 | 1518 | 8 | 6 | 7 | 0 | 0 | 12 | 0 | 0 | 0 | 31 | 58 | 12 |
| PD.36531 | 126 | 60 | 424 | 225 | 354 | 287 | 1 | 2 | 2 | 1 | 3 | 2 | 1 | 0 | 0 | 8 | 18 | 12 |
| PD.37301 | 778 | 814 | 848 | 1060 | 656 | 1369 | 0 | 0 | 0 | 0 | 0 | 0 | 0 | 0 | 0 | 0 | 0 | 0 |
| PD.36719 | 0 | 0 | 768 | 240 | 225 | 559 | 0 | 2 | 0 | 0 | 0 | 0 | 0 | 0 | 0 | 0 | 0 | 4 |
| PD.36347 | 613 | 1648 | 6669 | 2660 | 2601 | 1227 | 0 | 1 | 0 | 20 | 22 | 10 | 4 | 0 | 0 | 4 | 23 | 31 |
| PD.37795 | 6 | 175 | 1490 | 820 | 817 | 223 | 0 | 0 | 0 | 0 | 2 | 1 | 0 | 0 | 0 | 6 | 2 | 5 |
| PD.37701 | 867 | 4725 | 4811 | 868 | 221 | 1423 | 0 | 7 | 6 | 12 | 0 | 2 | 0 | 0 | 0 | 4 | 6 | 4 |
| PD.51830 | 427 | 2796 | 1282 | 2602 | 2500 | 3779 | 3 | 3 | 2 | 21 | 4 | 8 | 0 | 0 | 0 | 84 | 69 | 68 |
| PD.52568 | 68403 | 36188 | 178702 | 92859 | 99163 | 75564 | 1 | 0 | 0 | 3 | 3 | 2 | 0 | 0 | 8 | 3 | 9 | 8 |
| PD.51265 | 1773 | 207 | 4598 | 2927 | 3908 | 2564 | 1 | 2 | 0 | 2 | 2 | 8 | 0 | 0 | 0 | 3 | 8 | 5 |
| PD.52651 | 1189 | 1761 | 5445 | 6070 | 4931 | 4235 | 0 | 0 | 0 | 1 | 7 | 1 | 0 | 0 | 0 | 0 | 0 | 0 |
| PD.52670 | 922 | 2599 | 4537 | 5177 | 13669 | 15364 | 135 | 30 | 8 | 19 | 3 | 4 | 0 | 0 | 0 | 53 | 32 | 34 |
| PD.52649 | 152 | 236 | 1481 | 1717 | 317 | 912 | 0 | 2 | 0 | 3 | 1 | 2 | 0 | 0 | 0 | 4 | 4 | 5 |
| PD.52576 | 2339 | 2165 | 4008 | 34985 | 1332 | 13741 | 0 | 0 | 0 | 1 | 0 | 0 | 0 | 0 | 0 | 0 | 0 | 0 |
| PD.52359 | 40461 | 31448 | 164193 | 79981 | 83805 | 113442 | 23 | 981 | 93 | 470 | 458 | 453 | 457 | 78 | 100 | 357 | 701 | 361 |
| PD.70740 | 1775 | 1097 | 1622 | 2249 | 1446 | 4451 | 4 | 2 | 1 | 1 | 0 | 1 | 0 | 0 | 0 | 2 | 8 | 6 |
| PD.69752 | 5329 | 9593 | 8365 | 13355 | 13220 | 26101 | 11 | 25 | 9 | 72 | 50 | 23 | 4 | 2 | 2 | 7 | 66 | 0 |
| PD.69276 | 16175 | 4691 | 21005 | 31586 | 3812 | 23445 | 7 | 10 | 9 | 137 | 145 | 103 | 1 | 0 | 69 | 387 | 409 | 511 |
| PD.69863 | 1858 | 2133 | 22997 | 10654 | 10623 | 10927 | 8 | 4 | 24 | 47 | 37 | 82 | 2 | 11 | 1 | 30 | 46 | 24 |
| PD.70852 | 98 | 397 | 621 | 1070 | 557 | 1163 | 8 | 2 | 0 | 12 | 0 | 0 | 0 | 0 | 0 | 1 | 0 | 0 |
| PD.72277 | 12593 | 9579 | 18751 | 10383 | 16690 | 36684 | 52 | 42 | 9 | 31 | 8 | 2 | 0 | 0 | 0 | 1222 | 510 | 1141 |
| PD.72635 | 972 | 1153 | 1911 | 1059 | 1532 | 687 | 0 | 1 | 0 | 0 | 1 | 1 | 0 | 0 | 0 | 31 | 13 | 22 |
| PD.72244 | 237 | 498 | 1956 | 2896 | 113 | 516 | 14 | 5 | 0 | 6 | 4 | 6 | 0 | 0 | 0 | 9 | 1 | 1 |
| PD.73874 | 3272 | 6395 | 9377 | 6495 | 7496 | 11747 | 5 | 0 | 2 | 6 | 9 | 0 | 0 | 2 | 0 | 179 | 42 | 81 |
| PD.73932 | 4976 | 3805 | 9792 | 9615 | 7714 | 10531 | 22 | 20 | 12 | 28 | 39 | 35 | 0 | 0 | 0 | 228 | 129 | 126 |
| PD.73638 | 547 | 2618 | 6955 | 1968 | 9350 | 7981 | 0 | 1 | 0 | 23 | 57 | 35 | 0 | 0 | 0 | 25 | 18 | 7 |
| PD.73720 | 13637 | 7025 | 35278 | 21658 | 21449 | 7208 | 88 | 160 | 59 | 157 | 150 | 207 | 7 | 4 | 2 | 573 | 378 | 253 |
| PD.65627 | 237 | 3 | 798 | 326 | 518 | 457 | 0 | 0 | 0 | 0 | 0 | 0 | 0 | 0 | 0 | 0 | 0 | 0 |
| PD.66449 | 8828 | 2449 | 15735 | 9839 | 7933 | 2098 | 0 | 1 | 1 | 21 | 22 | 23 | 1 | 0 | 0 | 30 | 97 | 70 |
