## Supplementary Table 6 for "The genetic architecture of quantitative variation in the self-incompatibility response within *Phlox drummondii* (Polemoniaceae)"

| **QTL**  Table S6. Anther expressed candidate genes in QTL intervals. Gene IDs sourced from *Phlox drummondii* annotation. | **Linkage Group** | **5' CDS position** | ***P. drummondii* Annotation ID** | **TAIR ID** | **e-value** | **Identity (%)** | **Length (bp)** | **Arabidopsis Homolog Description** |
| --- | --- | --- | --- | --- | --- | --- | --- | --- |
| Q3A | 3 | 176927731 | MSTRG.56222 | No hit |  |  |  |  |
| Q3A | 3 | 311377118 | MSTRG.58220 | No hit |  |  |  |  |
| Q3A | 3 | 309189432 | MSTRG.58210 | AT1G24480 | 4e-81 | 72 | 582 | S-adenosyl-L-methionine-dependent methyltransferases superfamily protein |
| Q3A | 3 | 149173392 | MSTRG.55851 | AT3G44150 | 1e-17 | 66 | 433 | Expp1 protein |
| Q3A | 3 | 61208789 | MSTRG.59013 | AT5G15900 | 1e-48 | 65 | 1023 | RICHOME BIREFRINGENCE-LIKE, involved in the synthesis and deposition of secondary wall cellulose |
| Q3A | 3 | 242226686 | MSTRG.57286 | No hit |  |  |  |  |
| Q3A | 3 | 220256030 | MSTRG.56926 | AT4G38390 | 4e-115 | 69 | 1073 | root hair specific 17 |
| Q3A | 3 | 215217620 | MSTRG.60094 | AT5G28690 | 5e-22 | 76 | 143 | carboxylate clamp-TPR protein |
| Q3A | 3 | 230355123 | MSTRG.60173 | AT3G05610 | 4e-61 | 67 | 745 | Pectin methylesterase |
| Q3A | 3 | 149281454 | MSTRG.55856 | AT3G19020 | 4e-34 | 74 | 257 | LRX8, pollen expressed protein required for pollen tube growth |
| Q3A | 3 | 321335686 | MSTRG.58322 | No hit |  |  |  |  |
| Q3A | 3 | 6775100 | MSTRG.58543 | AT3G10360 | 1e-164 | 73 | 1036 | Arabidopsis Pumilio (APUM) proteins containing PUF domain, |
| Q3A | 3 | 235479358 | MSTRG.57123 | AT4G38190 | 0 | 74 | 3256 | encodes a gene similar to cellulose synthase |
| Q3A | 3 | 198902537 | MSTRG.56594 | AT2G16730 | 2e-18 | 65 | 567 | putative beta-galactosidase |
| Q3A | 3 | 162671255 | MSTRG.56035 | AT1G49270 | 3e-144 | 72 | 966 | proline-rich extensin-like  receptor kinase |
| Q3A | 3 | 145304306 | MSTRG.55789 | AT3G05120 | 2e-04 | 82 | 49 | gibberellin (GA) receptor ortholog of the rice GA receptor gene (OsGID1) |
| Q3A | 3 | 85988172 | MSTRG.59176 | AT3G46570 | 3e-16 | 72 | 137 | Glycosyl hydrolase superfamily protein |
| Q3A | 3 | 277375893 | MSTRG.57833 | No hit |  |  |  |  |
| Q3A | 3 | 176788570 | MSTRG.56219 | AT4G34940 | 2e-140 | 72 | 928 | Armadillo repeat protein, involved in the signaling network controlling tip growth and actin organization in the pollen tube |
| Q3A | 3 | 142659411 | MSTRG.55742 | AT3G19230 | 7e-32 | 64 | 1227 | malectin-like domain (MLD)-  and leucine-rich repeat (LRR)-containing protein 4 |
| Q3A | 3 | 147435425 | MSTRG.55831 | AT3G19090 | 5e-61 | 75 | 355 | Loss of function results in reduced transmission through male due to defects in pollen tube germination and pollen tube guidance |
| Q3A | 3 | 330583315 | MSTRG.58431 | AT2G39210 | 1e-99 | 72 | 718 | Major facilitator superfamily transmembrane transporter responsible for the uptake of picolinate herbicides |
| Q3A | 3 | 227127497 | MSTRG.56995 | AT4G38250 | 2e-69 | 66 | 1099 | Transmembrane amino acid transporter family protein |
| Q3A | 3 | 302661251 | MSTRG.58119 | AT3G18670 | 2e-06 | 73 | 95 | Ankyrin repeat family protein |
| Q3A | 3 | 328208511 | MSTRG.58410 | AT5G04390 | 5e-18 | 82 | 94 | C2H2-type zinc finger family protein |
| Q3A | 3 | 242923943 | MSTRG.57299 | AT3G04630 | 2e-07 | 71 | 112 | small gene family which have a KLEEK domain which may be involved in protein- protein interactions |
| Q3A | 3 | 51968332 | MSTRG.58936 | AT2G39380 | 3e-18 | 70 | 234 | member of EXO70 gene family, putative exocyst subunits, conserved in land plants |
| Q3A | 3 | 250576443 | MSTRG.57427 | AT5G39650 | 2e-58 | 70 | 552 | Sperm membrane protein that functions during fertilization to localize pollen membrane fusion factors to the pollen membrane in response to signals from the egg cell |
| Q3A | 3 | 159441052 | MSTRG.55998 | AT3G18830 | 2e-133 | 68 | 1440 | encodes a plasma membrane-localized polyol/cyclitol/monosaccharide-H+-symporter |
| Q3A | 3 | 302660036 | MSTRG.58118 | No hit |  |  |  |  |
| Q3A | 3 | 310462059 | MSTRG.58214 | No hit |  |  |  |  |
| Q3A | 3 | 213043463 | MSTRG.60078 | AT3G48640 | 0.008 | 83 | 42 | transmembrane protein |
| Q3A | 3 | 29245040 | MSTRG.58751 | AT3G52290 | 5e-33 | 66 | 643 | IQ67 (CaM binding) domain containing family |
| Q3A | 3 | 241790360 | MSTRG.60285 | No hit |  |  |  |  |
| Q3A | 3 | 131843812 | MSTRG.59492 | AT4G29210 | 6e-63 | 67 | 873 | gamma-glutamyltransferase (AKA gamma-glutamyl transpeptidase, EC 2.3.2.2) |
| Q3A | 3 | 27430989 | MSTRG.58729 | No hit |  |  |  |  |
| Q1A | 1 | 279819279 | MSTRG.37133 | AT1G77980 | 2e-37 | 75 | 230 | MIKC family of transcriptional regulators, pollen development and pollen tube growth |
| Q1A | 1 | 130977736 | MSTRG.34917 | AT5G35390 | 2e-64 | 69 | 637 | member of the receptor-like kinase family of genes. In pollen tubes, it accumulates in the plasma membrane of the apical growing tip |
| Q1A | 1 | 133555703 | MSTRG.34985 | No hit |  |  |  |  |
| Q1A | 1 | 307032085 | MSTRG.37568 | No hit |  |  |  |  |
| Q1A | 1 | 131876698 | MSTRG.34941 | AT2G24370 | 0 | 74 | 1121 | kinase with adenine nucleotide alpha hydrolases-like domain-containing protein |
| Q1A | 1 | 150486573 | MSTRG.35304 | AT5G01750 | 1e-05 | 67 | 163 | LURP-one-like protein |
| Q1A | 1 | 299197645 | MSTRG.37449 | AT3G12620 | 1e-141 | 72 | 963 | Protein phosphatase 2C family protein |
| Q1A | 1 | 284769004 | MSTRG.37209 | AT5G64790 | 3e-14 | 65 | 467 | O-glycosyl hydrolase |
| Q1A | 1 | 289749406 | MSTRG.37269 | No hit |  |  |  |  |
| Q1A | 1 | 246003121 | MSTRG.36570 | AT4G36220 | 7e-120 | 67 | 1434 | encodes ferulate 5-hydroxylase (F5H), involved in lignin biosynthesis |
| Q1A | 1 | 252575040 | MSTRG.36707 | AT2G45290 | 0.026 | 85 | 41 | Transketolase |
| Q1A | 1 | 241663487 | MSTRG.36511 | AT2G18180 | 1e-18 | 70 | 220 | Sec14p-like phosphatidylinositol transfer family protein |
| Q1A | 1 | 249690435 | MSTRG.36655 | No hit |  |  |  |  |
| Q1A | 1 | 284183064 | MSTRG.37195 | No hit |  |  |  |  |
| Q1A | 1 | 116310555 | MSTRG.34694 | No hit |  |  |  |  |
| Q1A | 1 | 306613315 | MSTRG.37559 | AT2G28320 | 3e-116 | 72 | 790 | Pleckstrin homology (PH) and lipid-binding START domains-containing protein |
| Q1A | 1 | 277802308 | MSTRG.37111 | AT5G10090 | 2e-69 | 66 | 1044 | Encodes one of the 36 carboxylate clamp (CC)-tetratricopeptide repeat (TPR) proteins |
| Q1A | 1 | 293027902 | MSTRG.37324 | AT5G13660 | 2e-15 | 68 | 305 | N-lysine methyltransferase |
| Q1A | 1 | 217292795 | MSTRG.36193 | No hit |  |  |  |  |
| Q1A | 1 | 307005967 | MSTRG.37564 | AT5G04220 | 1e-71 | 72 | 497 | protein specifically localized to the ER-PM boundary with similarity to synaptotagmins |
| Q1A | 1 | 178959534 | MSTRG.35730 | No hit |  |  |  |  |
| Q1A | 1 | 175591542 | MSTRG.35691 | ATCG01280 | 1e-152 | 92 | 378 | Chloroplast Ycf2 |
| Q1A | 1 | 159631213 | MSTRG.35447 | AT5G05700 | 5e-04 | 90 | 39 | arginyl-tRNA:protein transferase (ATE1), a component of the N-end rule pathway that targets protein degradation through the identity of the amino-terminal residue of specific substrates |
| S-locus & Q2C | 2 | 448262310 | MSTRG.52273 | AT5G63850 | 0.001 | 83 | 42 | Amino acid transporter whose expression is downregulated by dehydration |
| S-locus & Q2C | 2 | 419620539 | MSTRG.51556 | AT5G39200 | 1e-07 | 87 | 47 | myb-like protein Q |
| S-locus & Q2C | 2 | 445257081 | MSTRG.52161 | AT1G41880 | 0.015 | 92 | 26 | Ribosomal protein L35Ae family protein |
| S-locus & Q2C | 2 | 438900233 | MSTRG.51926 | AT5G39400 | 6e-32 | 70 | 1107 | Calcium/lipid-binding (CaLB) phosphatase |
| S-locus & Q2C | 2 | 443427665 | MSTRG.52078 | AT3G28150 | 3e-34 | 70 | 331 | TRICHOME BIREFRINGENCE-LIKE, shown to be involved in the synthesis and deposition of secondary wall cellulose |
| S-locus & Q2C | 2 | 429372178 | MSTRG.51744 | AT3G01270 | 6e-76 |  |  | Pectate lyase-like protein (PLL); enzymatic action of PLLs in the pollen grain, the pollen tube and the transmitting track contribute to an effective fertilization process |
| S-locus & Q2C | 2 | 448268403 | MSTRG.52274 | No hit |  |  |  |  |
| S-locus & Q2C | 2 | 445313134 | MSTRG.52164 | AT2G07040 | 3e-69 | 68 | 760 | Pollen receptor kinase. Coexpression of AtPRK2a with AtRopGEF12 resulted in isotropic pollen tube growth |
| S-locus & Q2C | 2 | 437556304 | MSTRG.51896 | AT1G74330 | 1e-49 | 72 | 1041 | Protein kinase superfamily protein |
| S-locus & Q2C | 2 | 445236483 | MSTRG.52158 | AT2G24370 | 3e-162 | 72 | 1064 | kinase with adenine nucleotide alpha hydrolases-like domain-containing protein |
| S-locus & Q2C | 2 | 408661476 | MSTRG.51375 | AT3G23570 | 3e-38 | 68 | 457 | alpha/beta-Hydrolases superfamily |
| S-locus & Q2C | 2 | 440745891 | MSTRG.51996 | AT3G55780 | 6e-11 | 75 | 112 | Glycosyl hydrolase superfamily protein |
| S-locus & Q2C | 2 | 454278109 | MSTRG.52597 | AT3G45400 | 1e-167 | 72 | 1101 | exostosin family protein |
| S-locus & Q2C | 2 | 445260776 | MSTRG.52162 | AT3G20200 | 2e-06 | 81 | 59 | kinase with adenine nucleotide alpha hydrolases-like domain-containing protein |
| S-locus & Q2C | 2 | 446605282 | MSTRG.52189 | AT2G19330 | 2e-126 | 71 | 978 | PIRL6, a distinct, plant-specific class of intracellular LRRs that likely mediate protein interactions, possibly in the context of signal transduction |
| S-locus & Q2C | 2 | 448276811 | MSTRG.52275 | No hit |  |  |  |  |
| S-locus & Q2C | 2 | 448262310 | MSTRG.52273 | AT5G63850 | 0.001 | 83 | 42 | Amino acid transporter whose expression is downregulated by dehydration |
| Q5A | 5 | 105226782 | MSTRG.70591 | AT1G20650 | 1e-140 | 73 | 879 | Protein kinase superfamily protein |
| Q5A | 5 | 81506051 | MSTRG.70301 | AT4G24390 | 0.02 | 88 | 34 | RNI-like superfamily protein. Auxin receptor F-box protein |
| Q5A | 5 | 23348962 | MSTRG.69322 | AT5G55040 | 0.013 | 87 | 38 | DNA-binding bromodomain-containing protein, interacts with core SWI/SNF complex components |
| Q5A | 5 | 103785283 | MSTRG.70562 | AT2G33670 | 1e-130 | 71 | 964 | homologs of the barley mildew resistance locus o (MLO) protein, acts redundantly with MLO9 to tether Ca2+ channels to the pollen tube plasma membrane to effect pollen tube guidance |
| Q5A | 5 | 57233391 | MSTRG.69882 | No hit |  |  |  |  |
| Q5A | 5 | 108261366 | MSTRG.70637 | AT1G76310 | 1e-127 | 73 | 809 | core cell cycle genes |
| Q5A | 5 | 22267069 | MSTRG.69280 | AT1G05577 | 5e-37 | 72 | 294 | SOK1, expressed during embryogenesis in the apical-lateral plasma membrane |
| Q5A | 5 | 44602140 | MSTRG.69669 | AT1G02040 | 5e-13 | 78 | 92 | C2H2-type zinc finger family protein |
| Q5A | 5 | 52713335 | MSTRG.69800 | AT2G20370 | 2e-44 | 66 | 845 | xyloglucan galactosyltransferase located in the membrane of Golgi stacks that is involved in the biosynthesis of fucose |
| Q5A | 5 | 41967380 | MSTRG.69619 | AT2G45010 | 2e-58 | 69 | 629 | PLAC8 family protein |
| Q5A | 5 | 140570225 | MSTRG.71067 | AT1G58215 | 1e-57 | 76 | 318 | spindle pole body component 110 |
| Q5A | 5 | 4376009 | MSTRG.68702 | AT3G56960 | 2e-44 | 76 | 254 | protein with phosphatidylinositol-4-phosphate 5-kinase activity that plays a role in pollen tip growth |
| Q5A | 5 | 12481522 | MSTRG.69002 | AT1G79400 | 9e-07 | 82 | 61 | member of Putative Na+/H+ antiporter family |
| Q5A | 5 | 15535121 | MSTRG.69098 | AT3G57690 | 6e-06 | 75 | 73 | putative arabinogalactan-protein (AGP23), recruited and transported by FH5 to maintain the tip growth of the pollen tube |
| Q5A | 5 | 148192194 | MSTRG.71172 | AT4G40030 | 3e-98 | 79 | 412 | Histone variant H3 |
| Q5A | 5 | 4377481 | MSTRG.68703 | AT3G07960 | 5e-121 | 75 | 674 | Encodes phosphatidylinositol-4-phosphate 5-kinase 6 (PIP5K6). Regulates clathrin-dependent endocytosis in pollen tubes |
| Q5A & Q5B | 5 | 347976885 | MSTRG.72547 | AT4G33390 | 1e-143 | 67 | 1691 | WEAK CHLOROPLAST MOVEMENT UNDER BLUE LIGHT-like protein |
| Q5A & Q5B | 5 | 363235864 | MSTRG.72677 | AT1G75160 | 2e-56 | 68 | 646 | BDR9 (BOUNDARY OF ROP DOMAIN9) |
| Q5A & Q5B | 5 | 381314894 | MSTRG.72903 | AT5G66310 | 2e-168 | 72 | 1078 | ATP binding microtubule motor family protein |
| Q5A & Q5B | 5 | 169098181 | MSTRG.71351 | AT5G20240 | 3e-76 | 74 | 486 | PISTILLATA, floral homeotic gene encoding a MADS domain transcription factor |
| Q5A & Q5B | 5 | 161042203 | MSTRG.71286 | No hit |  |  |  |  |
| Q5A & Q5B | 5 | 332125579 | MSTRG.72429 | AT5G02390 | 7e-08 | 71 | 129 | Target promoter of the male germline-specific transcription factor DUO1 |
| Q5A & Q5B | 5 | 314541838 | MSTRG.72312 | AT5G01610 | 2e-89 | 77 | 457 | hypothetical protein (Protein of unknown function, DUF538) |
| Q5B | 5 | 396615902 | MSTRG.73557 | No hit |  |  |  |  |
| Q5B | 5 | 386876573 | MSTRG.73031 | AT1G19940 | 8e-157 | 70 | 1377 | glycosyl hydrolase 9B5 |
| Q5B | 5 | 404716009 | MSTRG.74510 | No hit |  |  |  |  |
| Q5B | 5 | 405196615 | MSTRG.74578 | AT5G06380 | 6e-07 | 75 | 80 | hypothetical protein |
| Q5B | 5 | 404398182 | MSTRG.74455 | AT3G47440 | 6e-29 | 65 | 612 | functions as water and urea channels in pollen. Target promoter of the male germline-specific transcription factor DUO1 |
| Q5B | 5 | 390465907 | MSTRG.73204 | AT5G42340 | 4e-86 | 67 | 1122 | Plant U-box type E3 ubiquitin ligase |
| Q5B | 5 | 405514158 | MSTRG.74634 | No hit |  |  |  |  |
| Q5B | 5 | 404719505 | MSTRG.74512 | No hit |  |  |  |  |
| Q5B | 5 | 384074905 | MSTRG.72964 | AT1G75780 | 0 | 81 | 1322 | beta tubulin gene downregulated by phytochrome A (phyA)-mediated far-red light high-irradiance and the phytochrome B (phyB)-mediated red light high-irradiance responses |
| Q5B | 5 | 405276558 | MSTRG.74592 | AT3G54740 | 6e-38 | 74 | 246 | myosin |
| Q5B | 5 | 394970796 | MSTRG.73442 | AT5G42330 | 2e-09 | 81 | 68 | hypothetical protein |
| Q5B | 5 | 403326960 | MSTRG.74296 | No hit |  |  |  |  |
| Q5B | 5 | 404465199 | MSTRG.74472 | No hit |  |  |  |  |
| Q5B | 5 | 388386254 | MSTRG.73089 | AT1G75780 | 0 | 81 | 1322 | beta tubulin gene downregulated by phytochrome A (phyA)-mediated far-red light high-irradiance and the phytochrome B (phyB)-mediated red light high-irradiance responses |
| Q5B | 5 | 400858223 | MSTRG.73942 | AT2G13680 | 0 | 74 | 5685 | Responsible for the synthesis of callose deposited at the primary cell wall of meiocytes, tetrads and microspores. Required for exine formation during microgametogenesis and for pollen viability |
| Q5B | 5 | 395220819 | MSTRG.73473 | AT5G42340 | 9e-49 | 66 | 732 | Plant U-box type E3 ubiquitin ligase |
| Q5B | 5 | 404729532 | MSTRG.74514 | AT5G52340 | 0 | 74 | 1915 | member of EXO70 gene family, putative exocyst subunits |
| Q5B | 5 | 401792884 | MSTRG.74084 | No hit |  |  |  |  |
| Q5B | 5 | 400001569 | MSTRG.73861 | AT2G21720 | 3e-22 | 68 | 339 | ArgH (DUF639) |
| Q5B | 5 | 403148260 | MSTRG.74271 | AT1G08150 | 3e-04 | 80 | 51 | member of Putative Na+/H+ antiporter family |
| Q5B | 5 | 403169986 | MSTRG.74274 | No hit |  |  |  |  |
| Q5B | 5 | 404428380 | MSTRG.74464 | AT3G54700 | 0 | 74 | 1561 | member of the Pht1 family of phosphate transporters |
| Q5B | 5 | 401223139 | MSTRG.73996 | AT3G55470 | 1e-15 | 75 | 121 | Calcium-dependent lipid-binding (CaLB domain) family protein |
| Q5B | 5 | 397128140 | MSTRG.73605 | No hit |  |  |  |  |
| Q5B | 5 | 398537476 | MSTRG.73734 | AT1G22180 | 9e-66 | 71 | 574 | Sec14p-like phosphatidylinositol transfer family protein |
| Q4B | 4 | 260980391 | MSTRG.65780 | No hit |  |  |  |  |
| Q4B | 4 | 361268379 | MSTRG.66827 | No hit |  |  |  |  |
| Q4B | 4 | 289612897 | MSTRG.66102 | AT1G31810 | 1e-04 | 78 | 78 | Type II Arabidopsis formin14. Interacts with microtubules and microfilaments to regulate cell division |
| Q4B | 4 | 269418943 | MSTRG.65868 | AT4G17960 | 0.029 | 88 | 34 | phospholipid hydroperoxide glutathione peroxidase |
| Q4B | 4 | 182527164 | MSTRG.65158 | AT3G57530 | 2e-104 | 74 | 613 | Calcium-dependent Protein Kinase, ABA signaling component |
| Q4B | 4 | 147361361 | MSTRG.64803 | AT3G49540 | 0.001 | 84 | 44 | hypothetical protein |
| Q4B | 4 | 262895142 | MSTRG.65798 | AT3G15040 | 6e-15 | 70 | 188 | senescence regulator |
| Q4B | 4 | 175698847 | MSTRG.65101 | No hit |  |  |  |  |
| Q4B | 4 | 111052737 | MSTRG.64497 | AT1G17500 | 0 | 71 | 3438 | ATPase E1-E2 type family protein / haloacid dehalogenase-like hydrolase family protein |
| Q4B | 4 | 159053356 | MSTRG.64954 | No hit |  |  |  |  |
| Q4B | 4 | 158335242 | MSTRG.64936 | AT5G46910 | 1e-128 | 71 | 992 | H3K27me3 demethylase involved in temperature and photoperiod dependent repressing of flowering |
| Q4B | 4 | 111423234 | MSTRG.64503 | AT5G64570 | 2e-35 | 72 | 276 | ecreted beta-d-xylosidase that belongs to family 3 of glycoside hydrolases |
| Q4B | 4 | 358177203 | MSTRG.66764 | AT4G11990 | 0.011 | 79 | 48 | TPX2-LIKE Group A family with aurora binding andTPX2 domains |
| Q4B | 4 | 322514837 | MSTRG.66501 | AT5G52360 | 3e-79 | 76 | 398 | ADF10 is an actin-depolymerizing factor that preferentially binds ADP-G-actin and inhibits G-actin nucleotide exchange, regulates organization of actin filaments and vesicle trafficking during pollen tube growth |
| Q4B | 4 | 182762492 | MSTRG.65162 | AT1G04600 | 2e-44 | 69 | 459 | member of Myosin-like proteins |
| Q4B | 4 | 219133562 | MSTRG.65463 | No hit |  |  |  |  |
| Q4B | 4 | 210322896 | MSTRG.65373 | No hit |  |  |  |  |
| Q4B | 4 | 361170657 | MSTRG.66825 | No hit |  |  |  |  |
| Q4B | 4 | 311204924 | MSTRG.66351 | AT4G23500 | 3e-105 | 69 | 1008 | Pectin lyase-like superfamily protein |
| Q4B | 4 | 286631196 | MSTRG.66070 | No hit |  |  |  |  |
| Q4B | 4 | 209195366 | MSTRG.65370 | No hit |  |  |  |  |
| Q4B | 4 | 168486682 | MSTRG.65061 | No hit |  |  |  |  |

Note. TAIR ID is given for the *Arabidospis thaliana* homolog with the lowest E-Value following BLAST search in the TAIR database (arabidopsis.org)
