## Supplementary Table 7 for "The genetic architecture of quantitative variation in the self-incompatibility response within *Phlox drummondii* (Polemoniaceae)"

Table S7. Normalized count data for anther expressed candidate genes.

| ***P. drummondii* Annotation ID** | **Stigma and Style** | | | | | | **Anther** | | | **Leaf** | | | **Pollen** | | | **Stem** | | |
| --- | --- | --- | --- | --- | --- | --- | --- | --- | --- | --- | --- | --- | --- | --- | --- | --- | --- | --- |
|  | **1** | **2** | **3** | **4** | **5** | **6** | **1** | **2** | **3** | **1** | **2** | **3** | **1** | **2** | **3** | **1** | **2** | **3** |
| PD.56222 | 1 | 0 | 2 | 6 | 6 | 1 | 12365 | 1987 | 6241 | 4 | 2 | 2 | 4165 | 11104 | 14156 | 11 | 12 | 4 |
| PD.58220 | 0 | 0 | 0 | 13 | 7 | 1 | 6668 | 4037 | 4531 | 3 | 0 | 6 | 4986 | 7726 | 5925 | 7 | 2 | 8 |
| PD.58210 | 1 | 0 | 0 | 7 | 6 | 0 | 1418 | 984 | 925 | 3 | 0 | 3 | 1428 | 1680 | 1542 | 20 | 8 | 9 |
| PD.55851 | 0 | 4 | 2 | 11 | 0 | 3 | 1403 | 1321 | 1090 | 1 | 1 | 0 | 1920 | 1831 | 1645 | 11 | 0 | 11 |
| PD.59013 | 0 | 0 | 5 | 0 | 0 | 3 | 2430 | 1222 | 818 | 0 | 1 | 0 | 1933 | 1452 | 1270 | 2 | 1 | 1 |
| PD.57286 | 4 | 0 | 21 | 14 | 8 | 3 | 28602 | 20296 | 18267 | 9 | 7 | 26 | 24243 | 31385 | 35133 | 31 | 20 | 15 |
| PD.56926 | 1 | 3 | 2 | 23 | 0 | 8 | 3434 | 1933 | 2104 | 4 | 0 | 2 | 2838 | 3029 | 2757 | 28 | 8 | 9 |
| PD.60094 | 3 | 0 | 26 | 12 | 15 | 5 | 17565 | 12920 | 11153 | 4 | 2 | 20 | 17737 | 19776 | 19318 | 17 | 15 | 18 |
| PD.60173 | 107 | 136 | 152 | 352 | 336 | 313 | 74800 | 51077 | 48114 | 786 | 576 | 466 | 64512 | 76825 | 74663 | 414 | 241 | 317 |
| PD.55856 | 33 | 6 | 64 | 75 | 34 | 86 | 24049 | 21700 | 17370 | 30 | 53 | 22 | 23974 | 28289 | 33312 | 53 | 20 | 31 |
| PD.58322 | 5 | 0 | 13 | 21 | 38 | 31 | 40553 | 19369 | 21932 | 21 | 14 | 30 | 36558 | 38378 | 40574 | 52 | 19 | 30 |
| PD.58543 | 24 | 39 | 196 | 199 | 80 | 134 | 26276 | 17578 | 15450 | 118 | 56 | 76 | 22860 | 26145 | 28160 | 91 | 46 | 107 |
| PD.57123 | 66 | 71 | 384 | 335 | 177 | 324 | 12099 | 7935 | 8715 | 17 | 19 | 16 | 13080 | 13748 | 11546 | 35 | 11 | 26 |
| PD.56594 | 9 | 0 | 57 | 51 | 64 | 20 | 96131 | 61509 | 61371 | 46 | 15 | 70 | 79362 | 100142 | 95977 | 91 | 68 | 67 |
| PD.56035 | 15 | 17 | 57 | 163 | 157 | 197 | 23849 | 17710 | 16747 | 59 | 31 | 48 | 21959 | 27205 | 27796 | 83 | 61 | 52 |
| PD.55789 | 0 | 0 | 14 | 3 | 0 | 4 | 13389 | 12148 | 84 | 11 | 0 | 1 | 0 | 68 | 2312 | 19 | 4 | 10 |
| PD.59176 | 49 | 82 | 211 | 89 | 31 | 87 | 26785 | 19660 | 16940 | 25 | 50 | 33 | 27695 | 27472 | 22846 | 43 | 26 | 40 |
| PD.57833 | 0 | 0 | 0 | 0 | 0 | 3 | 2972 | 915 | 1468 | 0 | 0 | 5 | 3495 | 2551 | 462 | 4 | 0 | 2 |
| PD.56219 | 7 | 7 | 19 | 26 | 47 | 151 | 13173 | 6262 | 8966 | 11 | 8 | 10 | 13599 | 13955 | 11807 | 46 | 40 | 33 |
| PD.55742 | 37 | 39 | 84 | 510 | 140 | 118 | 21250 | 15703 | 15179 | 56 | 23 | 44 | 19286 | 25334 | 23776 | 39 | 33 | 117 |
| PD.55831 | 0 | 2 | 0 | 0 | 0 | 2 | 343 | 106 | 118 | 0 | 0 | 0 | 109 | 174 | 172 | 0 | 0 | 0 |
| PD.58431 | 10 | 8 | 176 | 88 | 156 | 106 | 7608 | 15058 | 4687 | 290 | 193 | 147 | 0 | 0 | 0 | 223 | 346 | 122 |
| PD.56995 | 3 | 1 | 5 | 0 | 1 | 6 | 26 | 36 | 28 | 0 | 0 | 0 | 48 | 31 | 166 | 0 | 2 | 0 |
| PD.58119 | 0 | 5 | 2 | 0 | 0 | 0 | 70 | 40 | 62 | 0 | 0 | 0 | 27 | 67 | 64 | 0 | 0 | 0 |
| PD.58410 | 3 | 2 | 4 | 46 | 81 | 7 | 2982 | 1851 | 1498 | 11 | 4 | 3 | 2944 | 2336 | 2638 | 43 | 17 | 40 |
| PD.57299 | 43 | 98 | 254 | 500 | 12 | 120 | 15445 | 10145 | 10969 | 39 | 21 | 13 | 13751 | 17197 | 14951 | 27 | 17 | 37 |
| PD.58936 | 0 | 0 | 13 | 0 | 2 | 46 | 3070 | 93 | 6957 | 113 | 36 | 43 | 0 | 0 | 0 | 20 | 7 | 7 |
| PD.57427 | 0 | 0 | 0 | 1 | 0 | 0 | 204 | 118 | 124 | 0 | 0 | 0 | 80 | 223 | 208 | 0 | 0 | 0 |
| PD.55998 | 0 | 0 | 0 | 0 | 0 | 0 | 135 | 58 | 60 | 0 | 0 | 1 | 57 | 112 | 95 | 0 | 0 | 0 |
| PD.58118 | 0 | 0 | 0 | 0 | 0 | 0 | 78 | 60 | 93 | 0 | 0 | 0 | 89 | 161 | 113 | 0 | 0 | 0 |
| PD.58214 | 0 | 0 | 0 | 0 | 0 | 0 | 2502 | 258 | 428 | 0 | 2 | 2 | 435 | 713 | 517 | 2 | 0 | 1 |
| PD.60078 | 3 | 5 | 0 | 85 | 4 | 0 | 125 | 830 | 181 | 14 | 0 | 0 | 76 | 325 | 85 | 0 | 0 | 16 |
| PD.58751 | 8 | 19 | 43 | 15 | 15 | 1322 | 21765 | 16718 | 15268 | 17 | 13 | 17 | 20849 | 24838 | 24849 | 41 | 29 | 34 |
| PD.60285 | 0 | 0 | 0 | 0 | 0 | 0 | 286 | 37 | 146 | 0 | 0 | 1 | 94 | 302 | 188 | 0 | 0 | 0 |
| PD.59492 | 0 | 0 | 0 | 0 | 0 | 0 | 43 | 43 | 17 | 0 | 0 | 0 | 24 | 30 | 48 | 0 | 0 | 1 |
| PD.58729 | 0 | 0 | 213 | 0 | 0 | 0 | 243 | 77 | 719 | 0 | 19 | 0 | 97 | 781 | 54 | 0 | 0 | 0 |
| PD.37133 | 1 | 0 | 5 | 4 | 1 | 5 | 6996 | 4269 | 4475 | 2 | 3 | 7 | 6230 | 7152 | 6653 | 4 | 10 | 13 |
| PD.34917 | 0 | 0 | 3 | 3 | 1 | 6 | 9491 | 6530 | 5863 | 4 | 2 | 5 | 10315 | 9306 | 8791 | 6 | 9 | 10 |
| PD.34985 | 0 | 0 | 0 | 2 | 2 | 3 | 6593 | 2125 | 2160 | 1 | 2 | 1 | 1609 | 4101 | 3554 | 1 | 3 | 2 |
| PD.37568 | 1 | 0 | 5 | 2 | 0 | 0 | 4526 | 2618 | 2506 | 2 | 0 | 3 | 2432 | 4273 | 2592 | 7 | 8 | 4 |
| PD.34941 | 3 | 1 | 19 | 10 | 15 | 7 | 22187 | 14994 | 17474 | 10 | 8 | 25 | 22148 | 29132 | 22075 | 16 | 17 | 14 |
| PD.35304 | 2 | 0 | 14 | 11 | 9 | 22 | 13021 | 10364 | 6566 | 7 | 2 | 2 | 10109 | 11434 | 11136 | 15 | 12 | 11 |
| PD.37449 | 65 | 78 | 188 | 233 | 179 | 217 | 8339 | 5690 | 5556 | 26 | 20 | 23 | 6862 | 8732 | 7585 | 23 | 41 | 40 |
| PD.37209 | 0 | 0 | 6 | 0 | 6 | 4 | 439 | 135 | 286 | 0 | 2 | 2 | 240 | 405 | 282 | 1 | 0 | 0 |
| PD.37269 | 0 | 2 | 4 | 32 | 14 | 6 | 5887 | 2615 | 3582 | 28 | 15 | 9 | 4245 | 5394 | 3464 | 23 | 14 | 32 |
| PD.36570 | 4 | 0 | 0 | 5 | 2 | 0 | 358 | 162 | 77 | 3 | 0 | 0 | 218 | 170 | 159 | 1 | 0 | 0 |
| PD.36707 | 12 | 0 | 27 | 50 | 0 | 18 | 59571 | 41467 | 40939 | 17 | 6 | 36 | 50764 | 66595 | 60923 | 67 | 37 | 51 |
| PD.36511 | 309 | 180 | 1325 | 1088 | 688 | 656 | 17014 | 11357 | 10826 | 41 | 64 | 64 | 12624 | 17390 | 17710 | 121 | 115 | 112 |
| PD.36655 | 0 | 0 | 3 | 0 | 0 | 0 | 637 | 530 | 652 | 0 | 0 | 1 | 473 | 1097 | 399 | 0 | 0 | 0 |
| PD.37195 | 0 | 0 | 0 | 2 | 1 | 0 | 923 | 486 | 229 | 0 | 0 | 1 | 633 | 419 | 396 | 2 | 0 | 0 |
| PD.34694 | 0 | 0 | 0 | 0 | 8 | 0 | 348 | 34 | 38 | 0 | 0 | 0 | 63 | 81 | 157 | 0 | 0 | 0 |
| PD.37559 | 11 | 2 | 9 | 108 | 217 | 60 | 10806 | 6213 | 5589 | 7 | 3 | 9 | 9291 | 9125 | 10634 | 15 | 6 | 8 |
| PD.37111 | 1 | 1 | 17 | 8 | 133 | 27 | 1566 | 892 | 811 | 43 | 14 | 8 | 1525 | 1213 | 1325 | 26 | 16 | 29 |
| PD.37324 | 0 | 0 | 0 | 0 | 2 | 0 | 92 | 69 | 161 | 0 | 1 | 0 | 61 | 223 | 106 | 3 | 1 | 0 |
| PD.36193 | 0 | 0 | 0 | 3 | 0 | 0 | 29 | 45 | 32 | 0 | 0 | 0 | 12 | 46 | 12 | 0 | 0 | 0 |
| PD.37564 | 0 | 0 | 0 | 0 | 0 | 0 | 92 | 61 | 61 | 0 | 0 | 0 | 61 | 95 | 91 | 0 | 0 | 3 |
| PD.35730 | 0 | 0 | 0 | 0 | 0 | 0 | 149 | 42 | 71 | 0 | 0 | 0 | 188 | 86 | 76 | 1 | 0 | 0 |
| PD.35691 | 0 | 0 | 0 | 0 | 0 | 0 | 61 | 42 | 31 | 0 | 0 | 0 | 9 | 88 | 104 | 0 | 0 | 0 |
| PD.35447 | 0 | 0 | 0 | 0 | 0 | 0 | 174 | 48 | 19 | 0 | 0 | 0 | 36 | 24 | 65 | 0 | 0 | 0 |
| PD.52273 | 5 | 1 | 9 | 16 | 9 | 8 | 28074 | 20190 | 17239 | 7 | 2 | 10 | 27382 | 29070 | 30481 | 36 | 15 | 22 |
| PD.51556 | 1 | 1 | 1 | 5 | 0 | 0 | 5433 | 3516 | 3360 | 0 | 1 | 3 | 4569 | 5293 | 5935 | 6 | 7 | 8 |
| PD.52161 | 0 | 0 | 0 | 11 | 0 | 9 | 2148 | 1279 | 917 | 1 | 0 | 0 | 1907 | 1589 | 2215 | 3 | 2 | 2 |
| PD.51926 | 41 | 50 | 59 | 167 | 105 | 163 | 6832 | 5277 | 5331 | 20 | 16 | 21 | 5728 | 8797 | 7390 | 20 | 10 | 15 |
| PD.52078 | 1 | 2 | 13 | 0 | 4 | 24 | 1648 | 1234 | 1743 | 0 | 0 | 0 | 1086 | 2753 | 1965 | 2 | 8 | 6 |
| PD.51744 | 25 | 0 | 78 | 69 | 44 | 67 | 123071 | 73438 | 70943 | 43 | 24 | 78 | 96469 | 113712 | 120643 | 123 | 86 | 87 |
| PD.52274 | 2 | 0 | 47 | 17 | 6 | 11 | 28646 | 13801 | 12881 | 13 | 3 | 13 | 18531 | 20638 | 29280 | 32 | 13 | 12 |
| PD.52164 | 225 | 140 | 214 | 287 | 498 | 134 | 75082 | 45448 | 42374 | 112 | 96 | 120 | 59075 | 73991 | 70462 | 81 | 146 | 76 |
| PD.51896 | 302 | 141 | 535 | 124 | 351 | 489 | 23161 | 17441 | 21207 | 38 | 56 | 61 | 22407 | 34501 | 31428 | 101 | 88 | 57 |
| PD.52158 | 1121 | 408 | 4602 | 2224 | 1879 | 1441 | 57890 | 42488 | 40580 | 88 | 91 | 105 | 53055 | 64875 | 61393 | 144 | 75 | 75 |
| PD.51375 | 0 | 0 | 2 | 0 | 0 | 0 | 776 | 383 | 516 | 0 | 0 | 0 | 385 | 824 | 1158 | 0 | 0 | 2 |
| PD.51996 | 0 | 0 | 0 | 1 | 0 | 0 | 1316 | 772 | 657 | 0 | 1 | 0 | 1177 | 1071 | 634 | 1 | 0 | 1 |
| PD.52597 | 0 | 0 | 0 | 0 | 2 | 0 | 629 | 151 | 342 | 0 | 0 | 0 | 89 | 565 | 243 | 0 | 1 | 1 |
| PD.52162 | 0 | 0 | 0 | 0 | 0 | 0 | 1060 | 505 | 488 | 0 | 0 | 1 | 713 | 705 | 1201 | 1 | 2 | 0 |
| PD.52189 | 2 | 4 | 9 | 41 | 54 | 153 | 1754 | 683 | 1259 | 1 | 1 | 2 | 1886 | 2156 | 1419 | 3 | 14 | 2 |
| PD.52275 | 32 | 0 | 404 | 42 | 13 | 42 | 49055 | 33779 | 29903 | 31 | 8 | 18 | 44842 | 50788 | 51386 | 57 | 38 | 35 |
| PD.52273.1 | 5 | 1 | 9 | 16 | 9 | 8 | 28074 | 20190 | 17239 | 7 | 2 | 10 | 27382 | 29070 | 30481 | 36 | 15 | 22 |
| PD.70591 | 1 | 4 | 7 | 16 | 8 | 3 | 11010 | 7357 | 6535 | 4 | 2 | 5 | 5259 | 11336 | 7470 | 14 | 7 | 12 |
| PD.70301 | 4 | 0 | 5 | 2 | 9 | 1 | 8594 | 5121 | 7083 | 3 | 7 | 3 | 6190 | 12707 | 13528 | 11 | 9 | 8 |
| PD.69322 | 2 | 2 | 2 | 17 | 10 | 12 | 8550 | 5682 | 5947 | 8 | 4 | 6 | 7367 | 10223 | 9460 | 13 | 6 | 7 |
| PD.70562 | 2 | 0 | 4 | 0 | 0 | 0 | 561 | 794 | 758 | 0 | 0 | 1 | 998 | 1188 | 875 | 0 | 0 | 0 |
| PD.69882 | 3 | 2 | 2 | 39 | 14 | 21 | 15615 | 9086 | 12654 | 19 | 5 | 14 | 15459 | 20632 | 23221 | 19 | 11 | 13 |
| PD.70637 | 1 | 3 | 8 | 34 | 12 | 0 | 3744 | 1697 | 1977 | 9 | 1 | 2 | 3038 | 3502 | 3173 | 63 | 82 | 76 |
| PD.69280 | 1 | 0 | 0 | 0 | 2 | 4 | 370 | 62 | 638 | 5 | 17 | 0 | 214 | 11 | 23 | 19 | 2 | 6 |
| PD.69669 | 3 | 0 | 0 | 0 | 0 | 8 | 515 | 166 | 71 | 0 | 0 | 0 | 118 | 120 | 111 | 2 | 0 | 0 |
| PD.69800 | 0 | 0 | 0 | 0 | 15 | 0 | 393 | 277 | 297 | 0 | 0 | 0 | 508 | 441 | 469 | 0 | 0 | 0 |
| PD.69619 | 0 | 1 | 0 | 2 | 0 | 0 | 786 | 304 | 592 | 0 | 1 | 0 | 681 | 843 | 911 | 4 | 1 | 0 |
| PD.71067 | 30 | 34 | 88 | 178 | 45 | 80 | 1137 | 598 | 599 | 35 | 31 | 5 | 977 | 948 | 1216 | 24 | 10 | 37 |
| PD.68702 | 0 | 0 | 0 | 5 | 4 | 18 | 438 | 31 | 212 | 5 | 0 | 2 | 214 | 361 | 267 | 4 | 4 | 0 |
| PD.69002 | 134 | 7 | 395 | 132 | 140 | 68 | 6994 | 6241 | 3893 | 94 | 112 | 70 | 6860 | 6363 | 7118 | 21 | 65 | 53 |
| PD.69098 | 1663 | 504 | 995 | 774 | 1142 | 603 | 47391 | 12786 | 14643 | 28 | 13 | 39 | 36973 | 24788 | 27550 | 425 | 187 | 291 |
| PD.71172 | 0 | 0 | 0 | 0 | 0 | 0 | 543 | 131 | 393 | 0 | 0 | 0 | 362 | 704 | 431 | 2 | 0 | 2 |
| PD.68703 | 0 | 0 | 0 | 0 | 3 | 107 | 813 | 349 | 418 | 0 | 11 | 5 | 399 | 631 | 554 | 7 | 0 | 2 |
| PD.72547 | 0 | 0 | 3 | 8 | 9 | 8 | 3036 | 1660 | 2105 | 0 | 0 | 4 | 1897 | 3087 | 2579 | 0 | 0 | 1 |
| PD.72677 | 3 | 0 | 31 | 2 | 8 | 8 | 9110 | 5113 | 5502 | 1 | 0 | 7 | 8362 | 9374 | 8897 | 5 | 8 | 9 |
| PD.72903 | 32 | 27 | 33 | 163 | 32 | 91 | 3618 | 7072 | 5191 | 56 | 8 | 4 | 3489 | 7861 | 7930 | 27 | 7 | 33 |
| PD.71351 | 0 | 3 | 10 | 1 | 21 | 39 | 684 | 1140 | 374 | 1 | 0 | 5 | 0 | 0 | 0 | 0 | 0 | 3 |
| PD.71286 | 0 | 5 | 0 | 3 | 0 | 0 | 134 | 103 | 30 | 2 | 0 | 0 | 267 | 52 | 33 | 1 | 0 | 0 |
| PD.72429 | 0 | 0 | 0 | 0 | 0 | 0 | 760 | 386 | 356 | 1 | 0 | 1 | 612 | 549 | 573 | 0 | 0 | 3 |
| PD.72312 | 24 | 0 | 0 | 3 | 0 | 0 | 185 | 29 | 47 | 0 | 0 | 0 | 28 | 67 | 60 | 0 | 0 | 0 |
| PD.73557 | 0 | 0 | 7 | 2 | 0 | 3 | 6970 | 3645 | 4229 | 3 | 0 | 7 | 5816 | 7751 | 5882 | 8 | 6 | 4 |
| PD.73031 | 6 | 0 | 3 | 4 | 9 | 6 | 9964 | 7114 | 5842 | 6 | 1 | 8 | 10223 | 9845 | 14353 | 6 | 2 | 7 |
| PD.74510 | 2 | 0 | 6 | 10 | 14 | 7 | 18103 | 12789 | 12526 | 7 | 3 | 10 | 18147 | 21474 | 19781 | 9 | 13 | 11 |
| PD.74578 | 4 | 1 | 18 | 16 | 9 | 11 | 23704 | 16043 | 19116 | 81 | 38 | 110 | 25302 | 33351 | 27209 | 35 | 25 | 25 |
| PD.74455 | 5 | 0 | 7 | 23 | 4 | 12 | 23610 | 17627 | 14930 | 7 | 9 | 15 | 29691 | 25771 | 23165 | 21 | 27 | 13 |
| PD.73204 | 12 | 7 | 5 | 9 | 10 | 17 | 17915 | 10610 | 12834 | 9 | 3 | 6 | 14989 | 20902 | 16457 | 13 | 20 | 13 |
| PD.74634 | 8 | 2 | 25 | 44 | 27 | 35 | 94532 | 46570 | 52953 | 32 | 6 | 77 | 73797 | 86463 | 67611 | 63 | 55 | 43 |
| PD.74512 | 0 | 0 | 27 | 17 | 13 | 1 | 24403 | 15032 | 17216 | 13 | 4 | 14 | 22165 | 30029 | 26418 | 33 | 22 | 23 |
| PD.72964 | 43 | 30 | 65 | 71 | 96 | 236 | 10051 | 6463 | 7989 | 33 | 16 | 35 | 10306 | 12766 | 10508 | 78 | 39 | 80 |
| PD.74592 | 4 | 2 | 55 | 23 | 21 | 8 | 21663 | 13237 | 12860 | 54 | 12 | 19 | 17282 | 22440 | 20288 | 44 | 27 | 27 |
| PD.73442 | 1 | 10 | 7 | 11 | 18 | 27 | 1840 | 461 | 965 | 3 | 3 | 2 | 1348 | 1476 | 1141 | 33 | 23 | 27 |
| PD.74296 | 9 | 12 | 25 | 17 | 23 | 16 | 484 | 1316 | 407 | 1 | 7 | 4 | 2 | 4 | 2 | 2 | 2 | 1 |
| PD.74472 | 2 | 6 | 22 | 26 | 5 | 21 | 857 | 261 | 327 | 0 | 6 | 1 | 489 | 560 | 507 | 1 | 0 | 5 |
| PD.73089 | 23 | 25 | 46 | 33 | 24 | 163 | 6902 | 3214 | 3191 | 13 | 12 | 10 | 7138 | 4820 | 5818 | 49 | 25 | 52 |
| PD.73942 | 162 | 20 | 124 | 201 | 132 | 196 | 28524 | 14545 | 16765 | 116 | 54 | 70 | 20828 | 24982 | 23529 | 99 | 62 | 109 |
| PD.73473 | 23 | 11 | 1 | 9 | 14 | 1 | 11644 | 7382 | 6321 | 10 | 2 | 13 | 9331 | 10936 | 11639 | 15 | 6 | 12 |
| PD.74514 | 0 | 11 | 0 | 3 | 0 | 7 | 939 | 240 | 421 | 0 | 0 | 1 | 930 | 634 | 694 | 7 | 3 | 3 |
| PD.74084 | 7 | 0 | 40 | 38 | 29 | 34 | 1027 | 640 | 666 | 7 | 7 | 3 | 1107 | 856 | 973 | 9 | 13 | 4 |
| PD.73861 | 0 | 0 | 0 | 5 | 0 | 0 | 294 | 186 | 105 | 1 | 0 | 0 | 218 | 181 | 264 | 0 | 0 | 2 |
| PD.74271 | 12 | 9 | 70 | 71 | 90 | 14 | 3538 | 2052 | 2986 | 41 | 21 | 17 | 2721 | 4966 | 3763 | 16 | 18 | 20 |
| PD.74274 | 283 | 55 | 542 | 826 | 126 | 485 | 37085 | 27340 | 26210 | 82 | 30 | 53 | 36819 | 43696 | 43127 | 53 | 49 | 64 |
| PD.74464 | 0 | 3 | 12 | 2 | 0 | 1 | 262 | 78 | 34 | 1 | 0 | 0 | 127 | 68 | 69 | 0 | 0 | 1 |
| PD.73996 | 21 | 8 | 1 | 11 | 31 | 13 | 319 | 355 | 513 | 1 | 0 | 0 | 374 | 811 | 393 | 7 | 1 | 1 |
| PD.73605 | 0 | 0 | 0 | 0 | 0 | 0 | 505 | 383 | 118 | 0 | 0 | 0 | 804 | 184 | 131 | 0 | 0 | 3 |
| PD.73734 | 0 | 0 | 0 | 0 | 0 | 0 | 547 | 56 | 143 | 0 | 0 | 3 | 73 | 232 | 179 | 0 | 0 | 0 |
| PD.65780 | 1 | 0 | 11 | 7 | 4 | 2 | 16505 | 12873 | 10520 | 5 | 0 | 12 | 15587 | 17626 | 17953 | 19 | 10 | 12 |
| PD.66827 | 0 | 0 | 0 | 5 | 3 | 8 | 12953 | 7100 | 5628 | 0 | 3 | 6 | 7282 | 8980 | 13447 | 16 | 8 | 5 |
| PD.66102 | 1 | 0 | 0 | 12 | 1 | 5 | 10948 | 9681 | 8272 | 6 | 1 | 11 | 9356 | 13422 | 13059 | 11 | 8 | 9 |
| PD.65868 | 2 | 0 | 15 | 17 | 28 | 10 | 35558 | 24851 | 21528 | 20 | 3 | 18 | 24644 | 39432 | 35800 | 35 | 28 | 20 |
| PD.65158 | 9 | 3 | 18 | 14 | 37 | 16 | 6513 | 2663 | 3749 | 5 | 2 | 4 | 5423 | 5799 | 4513 | 21 | 15 | 13 |
| PD.64803 | 3 | 0 | 7 | 15 | 4 | 42 | 12091 | 11066 | 11280 | 7 | 0 | 8 | 15041 | 18133 | 11146 | 21 | 10 | 8 |
| PD.65798 | 5 | 5 | 13 | 20 | 58 | 46 | 5327 | 5348 | 4197 | 124 | 46 | 136 | 4546 | 7042 | 8056 | 33 | 77 | 33 |
| PD.65101 | 17 | 1 | 44 | 26 | 36 | 50 | 1664 | 967 | 1075 | 2 | 3 | 4 | 1421 | 1787 | 1713 | 13 | 12 | 2 |
| PD.64497 | 328 | 198 | 1485 | 1504 | 467 | 776 | 49601 | 34680 | 31312 | 124 | 59 | 58 | 41625 | 50585 | 49861 | 104 | 105 | 227 |
| PD.64954 | 2 | 17 | 14 | 46 | 21 | 155 | 21034 | 16019 | 13582 | 15 | 6 | 16 | 19308 | 23512 | 28149 | 25 | 9 | 38 |
| PD.64936 | 2 | 10 | 13 | 10 | 15 | 2 | 505 | 129 | 169 | 8 | 8 | 3 | 172 | 295 | 257 | 0 | 5 | 7 |
| PD.64503 | 2 | 0 | 0 | 1 | 25 | 0 | 267 | 708 | 357 | 0 | 0 | 0 | 1081 | 525 | 690 | 1 | 2 | 0 |
| PD.66764 | 96 | 12 | 90 | 74 | 59 | 143 | 1826 | 1187 | 1012 | 7 | 17 | 1 | 1245 | 1708 | 1727 | 6 | 7 | 4 |
| PD.66501 | 22 | 38 | 139 | 516 | 95 | 227 | 3766 | 2007 | 2051 | 3 | 1 | 1 | 2480 | 3107 | 2707 | 8 | 1 | 1 |
| PD.65162 | 0 | 0 | 0 | 0 | 0 | 1 | 313 | 106 | 154 | 0 | 2 | 1 | 216 | 253 | 271 | 7 | 2 | 2 |
| PD.65463 | 0 | 0 | 0 | 1 | 0 | 0 | 281 | 118 | 109 | 0 | 0 | 0 | 76 | 178 | 155 | 0 | 0 | 0 |
| PD.65373 | 0 | 0 | 0 | 0 | 0 | 0 | 126 | 64 | 83 | 0 | 0 | 0 | 128 | 136 | 124 | 0 | 0 | 0 |
| PD.66825 | 0 | 0 | 0 | 0 | 0 | 0 | 130 | 585 | 397 | 0 | 0 | 0 | 4 | 716 | 352 | 0 | 1 | 0 |
| PD.66351 | 4 | 108 | 205 | 19 | 7 | 3 | 5233 | 1972 | 2382 | 5 | 4 | 1 | 3754 | 3484 | 4172 | 7 | 3 | 7 |
| PD.66070 | 0 | 0 | 0 | 0 | 0 | 0 | 43 | 120 | 113 | 0 | 0 | 0 | 316 | 88 | 28 | 0 | 1 | 0 |
| PD.65370 | 0 | 0 | 0 | 1 | 0 | 0 | 1366 | 56 | 63 | 5 | 18 | 0 | 74 | 109 | 113 | 2 | 0 | 1 |
| PD.65061 | 4 | 8 | 0 | 277 | 13 | 2 | 4544 | 842 | 35 | 0 | 0 | 1 | 0 | 0 | 0 | 0 | 0 | 0 |
